# Translational Quantitative Proteomic Assay for Bacteriophages: Opening a Frontier in Phage Pharmaceutical Development

**DOI:** 10.64898/2026.05.27.728049

**Authors:** Thomas D. Nguyen, Connor E. Gould, Jacob T. Sanborn, Joshua Tutin, Yan Pan, Haoran Gao, Donna Ruszaj, Devin Angevine, Jonathan Bussa, Daniel Atakora, Liang Chen, Dwayne R. Roach, Troy D. Wood, Nicholas M. Smith

**Affiliations:** Division of Clinical and Translational Therapeutics, School of Pharmacy & Pharmaceutical Sciences, University at Buffalo, Buffalo, New York, 14214, USA; Center for Integrated Global Biomedical Sciences, Department of Pharmacy Practice, School of Pharmacy & Pharmaceutical Sciences, University at Buffalo, Buffalo, New York, 14214, USA; Department of Pharmaceutical Sciences, School of Pharmacy & Pharmaceutical Sciences, University at Buffalo, Buffalo, New York, 14214, USA; Department of Biology, San Diego State University, San Diego, California, 92182, USA; Department of Chemistry, University at Buffalo, Buffalo, New York, 14260, USA

**Author notes:** Correspondence Nicholas M. Smith, PharmD, PhD, 308 Farber Hall, 360 Hayes Rd, Buffalo, New York, 14214.

**Keywords:** Bacteriophage therapy, liquid chromatography-tandem mass spectrometry, signature peptide, phage quantitation, pharmacokinetics, *Pseudomonas aeruginosa*

## Abstract

Bacteriophages are a promising therapeutic class against drug-resistant bacterial infections but require analytical methods meeting modern drug development standards. Plaque-based enumeration, the gold standard, is laborious, host-dependent, and highly variable. Targeted mass spectrometry of virion structural proteins offers an orthogonal, host-independent, scalable alternative. Here, we describe a targeted proteomic LC-MS/MS assay for the *Pseudomonas aeruginosa* podovirus LUZ19. Peptide mapping identified multiple structural proteins, providing 55% sequence coverage of a LUZ19 major capsid protein, from which the signature peptide EVAELDGQELAR was selected for quantitation. Using multiple reaction monitoring, calibrators and quality controls were evaluated across three independent runs, with calibrator bias of −7.0 to 4.6%, intra-day precision of 0.5 to 9.3%, and inter-day precision of 6.6 to 12.0%. Peptide concentrations were related to plaque-assay titers by weighted Deming regression, yielding an empirical copy number of 1,314 (95% CI: 1,098 to 1,529) peptide copies per PFU and a phage-equivalent concentration. This proof-of-concept study establishes the analytical foundation for a fully validated assay.

## Introduction

As antimicrobial resistance (**AMR**) continues to escalate into a global health crisis, conventional antibiotics are increasingly ineffective against a growing number of multidrug-resistant (**MDR**) and extensively drug-resistant (**XDR**) pathogens.^1, 2^ This alarming trend has reignited interest in bacteriophages (**phages**) as a therapeutic alternative to combat AMR.^3, 4^ Lytic phages, also known as virulent phages, are viruses that infect bacterial cells and replicate via the lytic cycle, wherein the infected bacterium is lysed to release newly formed phage progeny. However, the unique ability of phages to replicate in the presence of susceptible bacteria presents distinct challenges for both pharmacokinetic (**PK**) and pharmacodynamic (**PD**) analyses, particularly where distinguishing exogenously administered phages from endogenously replicated progeny phage can confound accurate quantitation and interpretation of phage exposure-response relationships during treatment.^5^

Currently, the double layer agar (**DLA**) assay remains the gold standard for phage enumeration, enabling quantitation of infectious phage particles within a solution.^6–8^ The DLA assay remains a valuable reference method, as it selectively quantifies phages capable of completing a full lytic cycle under defined conditions. While broadly applicable, the assay must be optimized for each specific phage-host pair, and results can be highly variable without rigorous standardization.^6, 9^ Even under standardized conditions, plaque enumeration is subject to significant variability arising from the Poisson distribution inherent to count data, establishing an irreducible analytical floor that persists independent of operator proficiency or procedural consistency.^9, 10^ While pragmatic for demonstrating therapeutic efficacy, this approach does not allow for the resolution of individual phage concentrations, which is critical for PK analyses that support cocktail design, dose selection, and identification of target concentrations. Moreover, the assay is labor-intensive, low-throughput, and susceptible to operator bias, making it impractical for high-throughput preclinical workflows or future clinical implementation.

Quantitative polymerase chain reaction (**qPCR**) assays have been developed for high-throughput phage enumeration by measuring genome copy number^9, 11, 12^ and have been used to measure kinetics in clinical phage therapy cases^13, 14^, supporting its proposed use as a scalable analytical approach for clinical trial implementation. While highly sensitive and adaptable to complex matrices, qPCR does not differentiate between infectious and non-infectious particles, and genome copy number does not reliably reflect functional titer, often resulting in overestimation of the biologically active phage population.^9, 12, 15^ In the context of PK/PD analyses, this distinction can be consequential, as phage therapy is governed by a dynamic predator-prey relationship in which phage replication, bactericidal activity, and resistance emergence are inextricably linked. Overestimated phage concentrations may therefore misguide dose selection, obscure true exposure-response relationships, and confound the pharmacodynamic indices necessary to characterize bactericidal activity over the course of treatment.^16^ This consideration applies equally to droplet digital PCR (**ddPCR**), which addresses amplification efficiency variability but shares the same scope limitation with respect to particle viability. Reported ddPCR-derived phage concentrations have been consistently higher than PFU counts, reinforcing that amplification-based methods reflect a complementary but distinct quantity from infectious particle concentration. Additionally, as phages replicate and undergo genetic drift over the course of treatment, mutations in the targeted genomic region may progressively affect quantitation accuracy, a consideration that is difficult to detect without orthogonal measurement.^17^ In a clinical setting where dosing decisions depend on accurate and reliable concentration measurements, the analytical scope of the measurement warrants careful consideration relative to the question being asked.

The translation of phage therapy into clinical practice hinges on the availability of robust analytical methods capable of supporting rigorous PK/PD characterization. Understanding exposure-response relationships is fundamental to phage candidate selection, dose selection, defining target concentrations, and characterizing the pharmacodynamic indices that link phage exposure to bactericidal activity and resistance suppression.^18^ These requirements demand analytical methods that are precise, accurate, reproducible, and capable of resolving individual phage concentrations across biological matrices. Neither current plaque assay nor nucleic acid-based methods fully meet these analytical demands. Liquid chromatography and tandem mass spectrometry (**LC-MS/MS**), as the established bioanalytical platform for drug development, is uniquely positioned to address these gaps by providing a standardized, matrix-flexible quantitation framework that can be developed in line with the regulatory expectations required for phage therapy clinical trial implementation.

LC-MS/MS using multiple reaction monitoring (**MRM**) offers high specificity, reproducibility, and high-throughput quantitation with multiplexing capabilities that are directly applicable to the demands of phage therapy development.^19, 20^ As the preferred platform for small molecule bioanalysis, LC-MS/MS operates within an established regulatory framework harmonized between the FDA and EMA under ICH M10, providing a defined path from initial proof of concept toward fully validated assays for clinical trial use. Despite these compelling advantages, its application in phage enumeration remains extremely limited.^21, 22^ To date, only a single study has applied mass spectrometry to develop a quantitative platform for a well-characterized filamentous phage.^23^ However, that study assessed only a single concentration and relied on prior knowledge of capsid protein copy numbers to infer phage particle counts. This approach would be challenging to extend to less well-characterized phages relevant to clinical application. The present study describes the development of a proteomic LC-MS/MS assay for the quantitation of a clinical *Pseudomonas aeruginosa* phage strain, LUZ19, leveraging conserved structural proteins as quantitative targets for host-independent phage quantitation (**Figure 1**). To assess the generalizability of this strategy across phylogenetically distinct phages, we further assessed its feasibility in another clinical *Pseudomonas aeruginosa* phage strain, E215, which diverges at the level of viral order, family, genus, and virion morphology.

**Figure 1:**
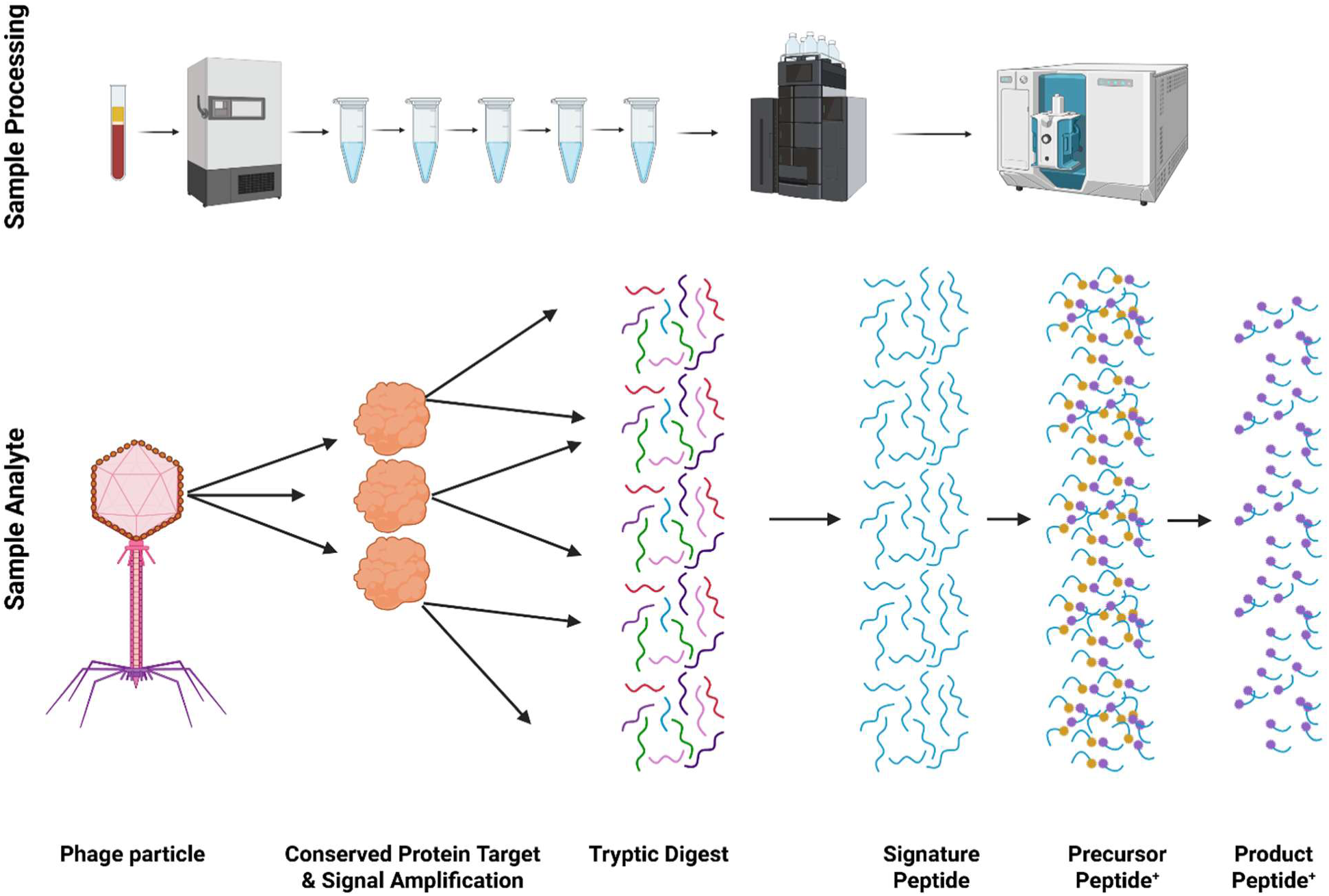
Conceptual overview of the targeted proteomic tandem mass spectrometry workflow for phage quantitation. Conserved structural proteins of the phage capsid serve as quantitative targets, leveraging conserved stoichiometry as an intrinsic signal amplification mechanism for reproducible, host-independent phage enumeration by tandem mass spectrometry (demonstrated here in phage lysates).

## Materials and Methods

### Materials and Reagents

Peptides **(Table S1**) were synthesized by Life Technologies Corporation (Carlsbad, CA, USA). For LUZ19, these were the signature peptide EVAELDGQELAR, derived from a LUZ19 major capsid protein, its light extended peptide EVAELDGQELARKFD (calibrator), and its stable isotope-labeled (SIL) extended peptide (internal standard). For E215, these were the signature peptide FITLGVEK, derived from an E215 major capsid protein, its light extended peptide QSKFITLGVEKRAK (calibrator), and its SIL extended peptide (internal standard). Purity was ≥95% for all peptides, and peptide content was determined by quantitative amino acid analysis (**AAA**). HPLC-grade acetonitrile, water, methanol, and acetone, as well as formic acid and acetic acid, were purchased from Fisher Scientific (Fair Lawn, NJ, USA). Trypsin from bovine pancreas, Type T (Sigma-Aldrich, St. Louis, MO, USA), sodium dodecyl sulfate (Thermo Scientific Chemicals, Waltham, MA, USA), iodoacetamide (Thermo Scientific Chemicals, Waltham, MA, USA), and dithiothreitol (Thermo Scientific, Rockford, IL, USA) were used.

### Production of Bacteriophage Stocks

*Pseudomonas aeruginosa* laboratory strain PAO1 was cultured in cation adjusted Mueller-Hinton Broth (Mg^2+^ 25 mg/L and Ca^2+^ 12.5 mg/L) or agar (Becton Dickinson, Sparks, MD, USA) at 37°C. Podovirus LUZ19, classified within the genus *Phikmvvirus*, and myovirus E215, classified within the genus *Pbunavirus*, were isolated on PAO1. Phage propagation and purification were performed as previously described.^8^ Briefly, phage lysates were sterilized by 2x high-speed centrifugation and 0.2 µm dead-end filtration.

### Double Layer Agar Assay for Bacteriophage Titration

Phages were quantified by double-aliquot 48-spot serial titration, as previously described.^8^ Briefly, PAO1 cultured at OD_600_ 0.2 – 0.3 was lawned and dried over agar medium. Two identical 8-well columns of tenfold serially diluted phage samples were spotted at 4 µL, in sextuplicate. Titrations were incubated at 37°C until plaque forming units (**PFU**) were visible. Titer was calculated per spot (Equation 1) using discrete, countable plaques. Spots with no plaques, and spots with overlapping plaques that prevented reliable counting, were excluded. The detection limit was 250 PFU/mL. Titers are reported as the geometric mean of technical sextuplicate.

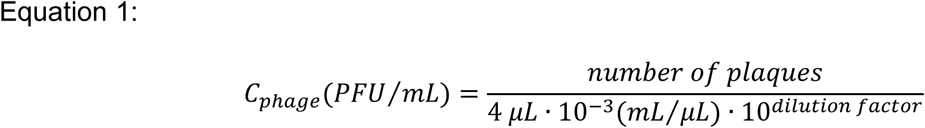

### Sample Preparation

A modified surfactant-aided-precipitation/on-pellet-digestion (**SOD**) strategy was employed for sample cleanup.^24^ A 285 μL aliquot of phage lysate per sample was prepared in a solution containing 1.0% sodium dodecyl sulfate and incubated at 90°C for 10 minutes to denature phage capsids. Dithiothreitol (**DTT**) was added to a final concentration of 9.8 mM, followed by incubation at 56°C for 30 minutes to reduce proteins; then iodoacetamide (**IAM**) was added to a final concentration of 58.5 mM and incubated in the dark for 30 minutes at 37°C to prevent reformation of disulfide bonds. One volume of chilled acetone was added to ensure no visible particulate was observed; another three volumes of chilled acetone were then added, followed by 3 hours incubation at −20°C. Post-incubation, samples were centrifuged at 30,279 g at 4°C for 30 minutes, then the supernatant was poured out, washed with one volume of chilled acetone, and the pellet was allowed to air-dry. The pellet was reconstituted in ammonium bicarbonate (50 mM, pH = 8.3) and the SIL internal standard was added to a final concentration of 1.2 ng/mL, followed by 2.5 µg of trypsin, to a final volume of 100 μL. Samples were incubated overnight at 37°C to achieve complete digestion. Digestion was terminated with formic acid at a final concentration of 0.5%.

### Signature Peptide Selection and Identification

Peptides derived from phage structural proteins were separated on an Acclaim PepMap 100 (75 µm x 15 cm, 2 µm) nanoViper C18 analytical column on an EASY-nLC 1200 system (Thermo Fisher Scientific, Waltham, MA, USA). Separation was achieved through a gradient elution method with mobile phases consisting of 0.1% formic acid in water (Mobile Phase A) and 0.1% formic acid in acetonitrile (Mobile Phase B) at a flow rate of 300 nL/min. The following gradient was applied: 0% B to 2% B at 10 min, 2% B to 40% B at 45 min, 40% B to 100% B at 50 min, followed by a 5-min wash at 100% B. Separated peptides were analyzed using a LTQ Orbitrap XL system (Thermo Fisher Scientific, Waltham, MA, USA). The samples were analyzed in positive mode using the following tune parameters: source voltage 1.5kV, capillary voltage 35V, tube lens 100V. Tandem mass spectrometry was performed using untargeted data-dependent MS^2^ with the top 10 highest abundant precursors selected for fragmentation. Peptide mapping of structural proteins for LUZ19 and E215 was performed using the Sequest search algorithm in Proteome Discoverer (version 2.3, Thermo Fisher Scientific, Waltham, MA, USA) and matched against the NCBI FASTA database. To assess specificity, the signature peptides were additionally evaluated against the *Pseudomonas aeruginosa* PAO1 proteome. Signature peptides were selected from high-abundance structural capsid proteins based on peptide specificity, signal intensity, and chromatographic performance.

### LC-MRM-MS Analysis

Quantitative analysis was performed on an analytical flow LC system coupled to a triple-quadrupole MS. An ExionLC AE was interfaced to a QTRAP 7500+ mass spectrometer (SCIEX, Framingham, MA, USA) via an ESI source. LUZ19 peptides were separated on an XSelect Premier HSS T3 column (2.1 mm × 150 mm, 3.5 μm, Waters, Milford, MA, USA). A 15-minute gradient was employed with a flow rate of 0.200 mL/min. Injection volumes ranged from 5 to 25 μL, with larger volumes used at lower concentrations. Mobile phases consisted of 10% methanol in water containing 0.1% formic acid (Mobile Phase A) and 95% acetonitrile in water containing 0.1% formic acid (Mobile Phase B). The gradient was initiated at 14% B, increasing to 17% B at 6.5 min, then to 23% B at 8 min, followed by a step to 95% B from 8.1 to 10.5 min, and re-equilibrated at 14% B from 10.6 to 15 min. The column temperature was 40°C. The spray voltage was 2.5 kV, and the source temperature was 415°C. Quantitation of LUZ19 was performed using multiple reaction monitoring of the transitions *m/z* 665.3 → 673.5 [(M+2H)^2+^ → y_6_, monitored], *m/z* 665.3 → 788.5 [(M+2H)^2+^ → y_7_, quantifier], and *m/z* 665.3 → 901.5 [(M+2H)^2+^ → y_8_, qualifier] for the signature peptide, and *m/z* 670.5 → 798.2 [(M+2H)^2+^ → y_7_, quantifier], and *m/z* 670.5 → 1040.4 [(M+2H)^2+^ → y_9_, qualifier] for the SIL internal standard (**Table S2**). The E215 method and multiplex method used for mixed LUZ19 and E215 samples are described in the **Supporting Information**. MRM transitions and collision energies for each peptide were optimized by direct analysis of the signature peptide and detectability in phage lysate was confirmed prior to calibration. For LUZ19, the optimized collision energy was set to 32 V, the Q0 dissociation was set to 70 V, and dwell time was 150 ms for each transition. Data acquisition and processing were performed using SCIEX OS (version 3.3.0, SCIEX, Framingham, MA, USA).

### Preparation of Calibrators and Establishment of Calibration

Calibrator solutions of the light extended peptide EVAELDGQELARKFD were prepared in 50 mM ammonium bicarbonate at 0.004, 0.008, 0.08, 0.8, 8, 50, and 80 ng/mL and underwent tryptic digestion only.^25^ The SIL extended peptide, incorporating [^13^C_6_, ^15^N_4_]-arginine, was used as the internal standard (**IS**) at a fixed concentration of 1.2 ng/mL for LUZ19 single-phage samples and 1 ng/mL for each IS in the LUZ19-E215 mixtures. The IS was added to calibrators, QCs, and phage lysate samples immediately before digestion, correcting for variability in tryptic digestion and instrument response. Quantitation was performed using peak area ratios of the signature peptide relative to the SIL internal standard. Analyte identity was assessed by the qualifier-to-quantifier peak area ratio and ratios within ± 20% were considered consistent. Linear regression with a 1/*x*^2^ weighting factor was applied to construct the calibration curve. The peptide LLOQ was defined as the lowest calibration level and its phage-equivalent concentration was calculated using Equation 4.

### Determination of the Empirical Copy Number

Three independent phage lysate stocks were prepared by 10-fold and half-log serial dilutions over a concentration range of 10^5^ to 10^9^ PFU/mL using the SOD strategy. Peak area ratios of the signature peptide released from digested phage lysate samples were plotted against a calibration curve to derive peptide concentrations, which were subsequently converted to femtomolar units (**Equation 2**). Phage-equivalent concentration is defined here as the particle concentration that would produce the observed peptide signal at the empirical copy number. It is not interchangeable with PFU/mL, which counts only particles able to complete a productive infection on the titration host. The empirical copy number is defined as the number of peptide copies detected per PFU, reflecting the copy number of the protein per virion together with the ratio of total particles to PFU and the efficiency of capsid disruption, digestion, peptide recovery, and sample processing. Nominal phage concentrations were independently converted from PFU/mL to femtomolar units using Avogadro’s number (**Equation 3**), with the caveat that PFU/mL quantifies infectious particles capable of completing a lytic cycle on the host isolate and may underestimate total virion concentration. Weighted Deming regression was performed in R to relate peptide molarity to DLA-derived particle molarity, with the slope of the regression used as the empirical copy number. Digested E215 lysate was similarly prepared by serial dilution across 10^5^ – 10^10^ PFU/mL and peptide molarity was converted to DLA-derived particle molarity to estimate the empirical copy number.

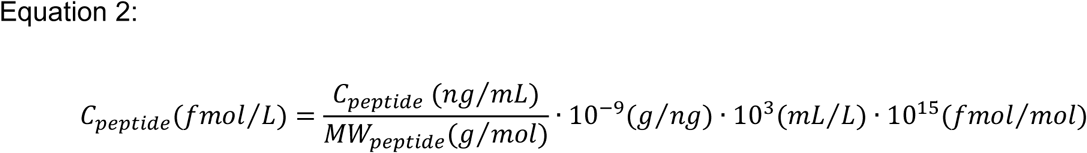

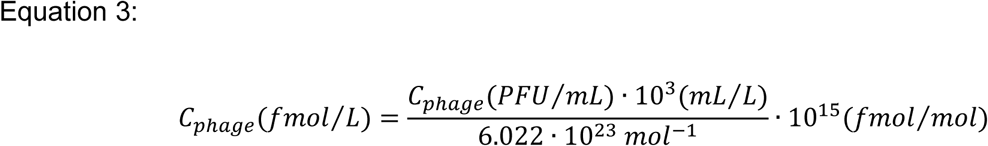

### Statistical Analyses

Calibration curves were constructed by linear regression with a 1/x^2^ weighting factor, selected because response variability scaled with concentration across the calibration range. The empirical copy number was estimated by weighted Deming regression of peptide molarity against DLA-derived particle molarity, accounting for measurement error in both analytical methods. The error ratio (λ) was estimated as the pooled variance of log-transformed peptide area ratios from replicate QC injections divided by the pooled variance of log-transformed plaque titers, scaled by the number of titer replicates averaged per lysate, and the regression slope and intercept are reported with 95% confidence intervals. Each lysate contributed one point, using the geometric mean of its plaque titers. Lysates were included if back-calculated peptide concentration fell within the calibration range and if the peptide concentration was within ± 20% of that predicted from the 10^8^ PFU/mL lysate of the same stock by the nominal dilution. Sensitivity of the empirical copy number to the concentration range and stock-specific estimates are shown in **Supporting Information**. For each stock, peak area ratio was regressed on DLA titer by weighted linear regression (1/x^2^ weighting) of untransformed data and proportionality was assessed from the slope of log_10_-transformed data. The double layer agar assay was performed in sextuplicate at each dilution level across three independent phage stocks (A, B, and C). LC-MS/MS calibration standards were analyzed once per run (n = 7) and quality control samples were assessed in triplicate across three independent analytical runs performed on separate days for a total n = 9 per quality control level for inter-day precision and accuracy. For QC lysates, intra- and inter-day precision of predicted phage-equivalent concentrations was expressed as the geometric CV, calculated from the standard deviation of the log-transformed concentrations within day (n = 3) and across days (n = 9). Agreement between LC-MS/MS-derived phage-equivalent concentrations and double layer agar-derived titers was assessed from the paired difference of log_10_-transformed concentrations (MS-derived minus DLA-derived). Each QC lysate was titered once and its geometric mean DLA titer (n = 6 replicates) served as the common reference for all MS measurements of that lysate. MS replicates were averaged within day (n = 3 replicates). Bias was calculated as the mean of these paired differences (one per lysate level per day) and 95% limits of agreement were calculated as the bias ± 1.96 times the combined standard deviation of MS between-day and DLA replicate variability. Results are reported as fold differences, with 95% confidence intervals that include both MS between-day variability and DLA replicate uncertainty (Welch-Satterthwaite degrees of freedom). The bias, its 95% confidence interval, and limits of agreement were compared with the 0.33 log_10_ (2.14-fold) threshold proposed for agreement between DLA replicates.^9^ In LUZ19-E215 mixtures, measured phage-equivalent concentrations and DLA titers were compared with expected values as the mean log_10_ difference. Cross-phage interference was assessed separately for each phage by linear regression of the log_10_ phage-equivalent concentration of each sample preparation on the log_10_ expected titer of the same phage and of the co-occurring phage and was evaluated from the coefficient of the co-occurring phage term (α = 0.05). Statistical analyses were performed in R (version 4.4.2, R Foundation for Statistical Computing, Vienna, Austria).

### Phage Directed Evolution and Host Range Expansion

Nine clinical *Pseudomonas aeruginosa* isolates (AR-0236, AR-0240, AR-0245, AR-0246, AR-0250, AR-0254, AR-0257, AR-0259, and AR-0270) obtained from the CDC/FDA Antibiotic Resistance (**AR**) Isolate Bank were used for directed evolution and efficiency of plating (**EOP**) assessment. Directed evolution of LUZ19 was performed using a modified single-phage Appelmans protocol.^26^ Using a 96-well plate format, serial 10-fold dilutions of LUZ19 were prepared across each row, with each row corresponding to a distinct clinical isolate. Following overnight incubation at 37°C, wells exhibiting clearing and the first turbid transition well were pooled across all rows into a single master lysate, clarified by centrifugation, and filter-sterilized. This evolved phage lysate was then used as the input for the subsequent rounds of selection, with the process repeated for 10 rounds to promote host range expansion. EOP was calculated as the ratio of PFU/mL obtained on each clinical isolate relative to the PFU/mL obtained on the reference strain PAO1, assessed at Day 0 and following directed evolution using the DLA method.^27^ Each EOP was determined in three independent experiments, each with three technical plaque replicates, and is reported as mean ± SD of the three independent experiments. EOP was classified as high (≥ 0.5), medium (0.1 to < 0.5), low (> 0.0001 to < 0.1), or inefficient (≤ 0.0001), and host-range expansion was defined as an increase of at least one class after directed evolution^28^.

### Evaluation of Quantitative Performance

Method performance was evaluated using three independent analytical runs on separate days to assess intra- and inter-day variability. Acceptance criteria from ICH M10 were applied as benchmarks of reproducibility; a full ICH M10 validation was not performed. Quality control (**QC**) samples of the light extended peptide EVAELDGQELARKFD, prepared at 0.006, 0.4, and 40 ng/mL, were evaluated alongside each calibration curve to assess accuracy and precision across runs. Selectivity was evaluated by analyzing blank cation-adjusted Mueller-Hinton broth carried through sample preparation and uninfected PAO1 culture lysate. To evaluate the accuracy of the empirical copy number, phage lysate QC samples were prepared from both unmodified LUZ19 and evolved LUZ19 at approximately 10^5^, 10^7^, and 10^9^ PFU/mL. Aliquots of these QC samples were assessed in triplicate for each of the three different days. The empirical copy number derived from stocks A, B, and C was fixed before evaluation. Phage-equivalent concentrations estimated from the MS method using the empirical copy number (**Equation 4**) were compared to DLA-derived titers of QC lysates prepared from stocks independent of those used to derive it. Lysates at ∼10^5^ PFU/mL, which produced peptide responses below the LLOQ, were excluded from the agreement analysis.

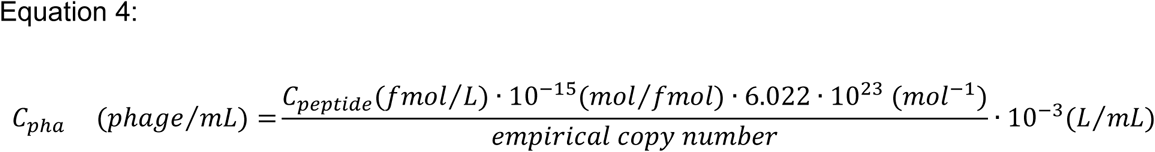

### Data availability

Mass spectrometry data have been deposited in the MassIVE repository under accession number MSV000101576 (ProteomeXchange identifier PXD077596) and are publicly accessible at <u>doi:10.25345/C5RX93T21</u>. All other data supporting the findings of this study are included in the article or are available from the corresponding author upon reasonable request.

## Results

### Proteomic Characterization and Signature Peptide Selection

To prioritize candidates for quantitative method development, proteins were filtered based on structural annotation and abundance, resulting in four proteins identified from the NCBI FASTA database for further evaluation (**Table 1**). The identified proteins included major capsid and minor capsid proteins along with two hypothetical proteins. The major capsid protein yielded 55% sequence coverage from 15 identified peptides. The remaining proteins featured varying coverage coming from large tryptic peptides. The two hypothetical proteins YP_001671995.1 and YP_001671965.1 featured 46% and 2% sequence coverage derived from two and one tryptic peptides, respectively. Due to the high degree of sequence coverage and number of peptides identified, peptides derived from the major capsid protein YP_001671977.1 were identified as potential candidate signature peptides. The signature peptide EVAELDGQELAR was selected as the quantitative peptide for the major capsid protein YP_001671977.1, identified with 8.72 ppm mass error for the precursor and an MS^2^ spectrum (**Figure 2**). The peptide was selected based on appropriate size for quantitation, high signal intensity post-digestion, and absence of easily modifiable amino acids. MRM transitions for EVAELDGQELAR were subsequently confirmed in both the calibration standards and the digested LUZ19 phage lysate, demonstrating consistent detectability and retention times (**Figure 3A**). EVAELDGQELAR did not match any of the 5,560 protein sequences in the PAO1 proteome. The ion ratio was stable across the calibration range (mean 0.74, CV 4.1%), and calibrators, QCs, and quantifiable lysate injections were within ± 20% of the calibrator mean (**Table S4**). A calibration curve for the synthetic signature peptide (undigested) demonstrated a linear response across a concentration range of 0.04 to 8 ng/mL (**Figure 3B and S1A**), confirming the intrinsic proportionality of the peptide prior to digestion. The calibration curve of the light extended peptide EVAELDGQELARKFD, which is digested to release the signature peptide and used to quantify digested phage samples, demonstrated a linear response across the concentration range of 0.004 to 80 ng/mL with back-calculated bias within ± 11.0% (**Figure 3C**).

**Figure 2:**
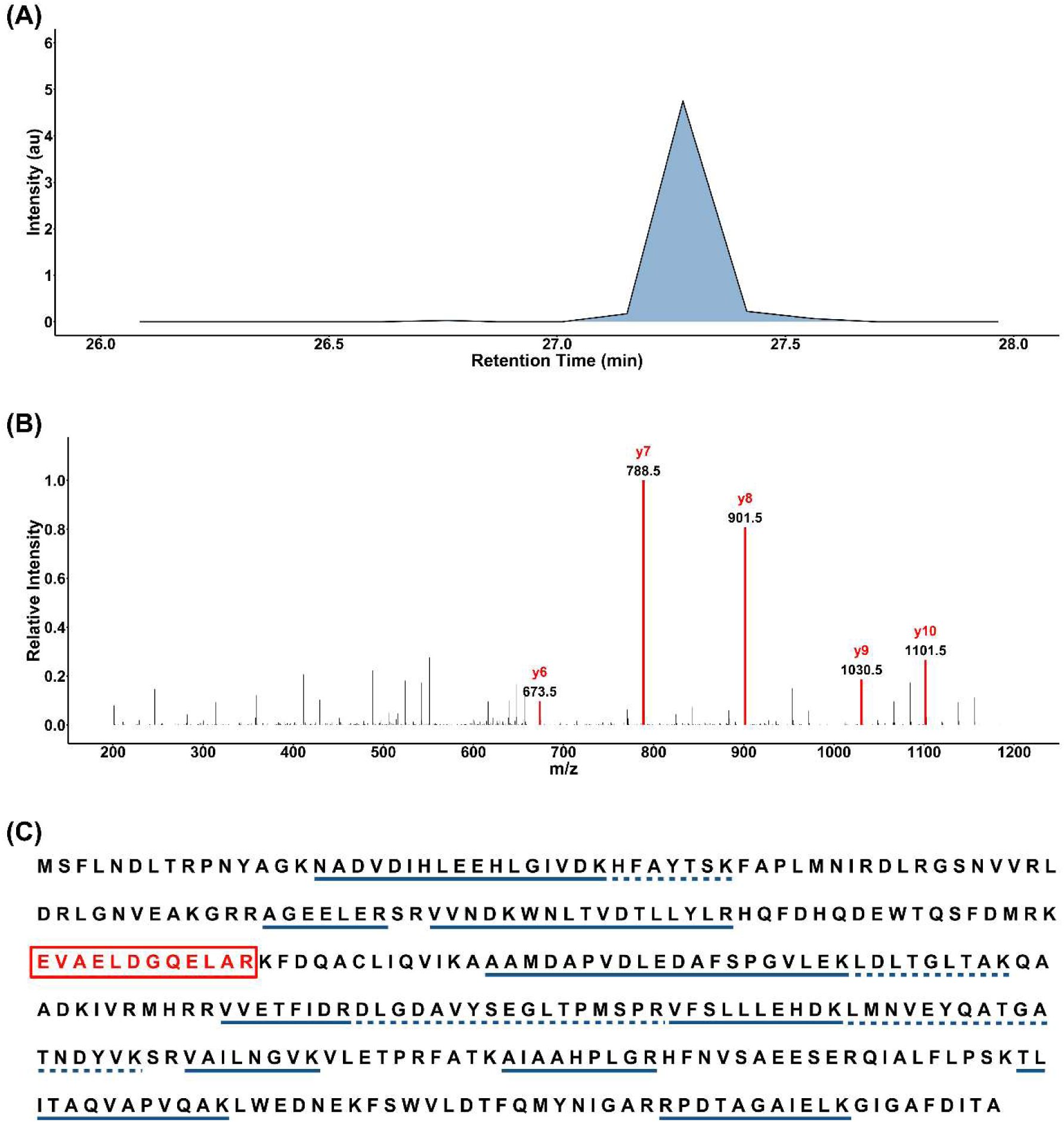
Discovery-phase identification of LUZ19 signature peptide EVAELDGQELAR by high-resolution Orbitrap mass spectrometry. (A) Extracted ion chromatogram of the precursor ion corresponding to EVAELDGQELAR (*m/z* = 665.3414), showing a single chromatographic peak at a retention time of ∼27.3 min. (B) MS^2^ fragmentation spectrum of the precursor ion, with y-ion fragments used for peptide identification and subsequent MRM transition selection highlighted in red (y6 = 673.5, y7 = 788.5, y8 = 901.5, y9 = 1030.5, and y10 = 1101.5 *m/z*). (C) Sequence coverage map of the LUZ19 major capsid protein with all identified tryptic peptides underlined and EVAELDGQELAR (residues 113-124) boxed in red to indicate its selection as the primary quantitative signature peptide.

**Figure 3:**
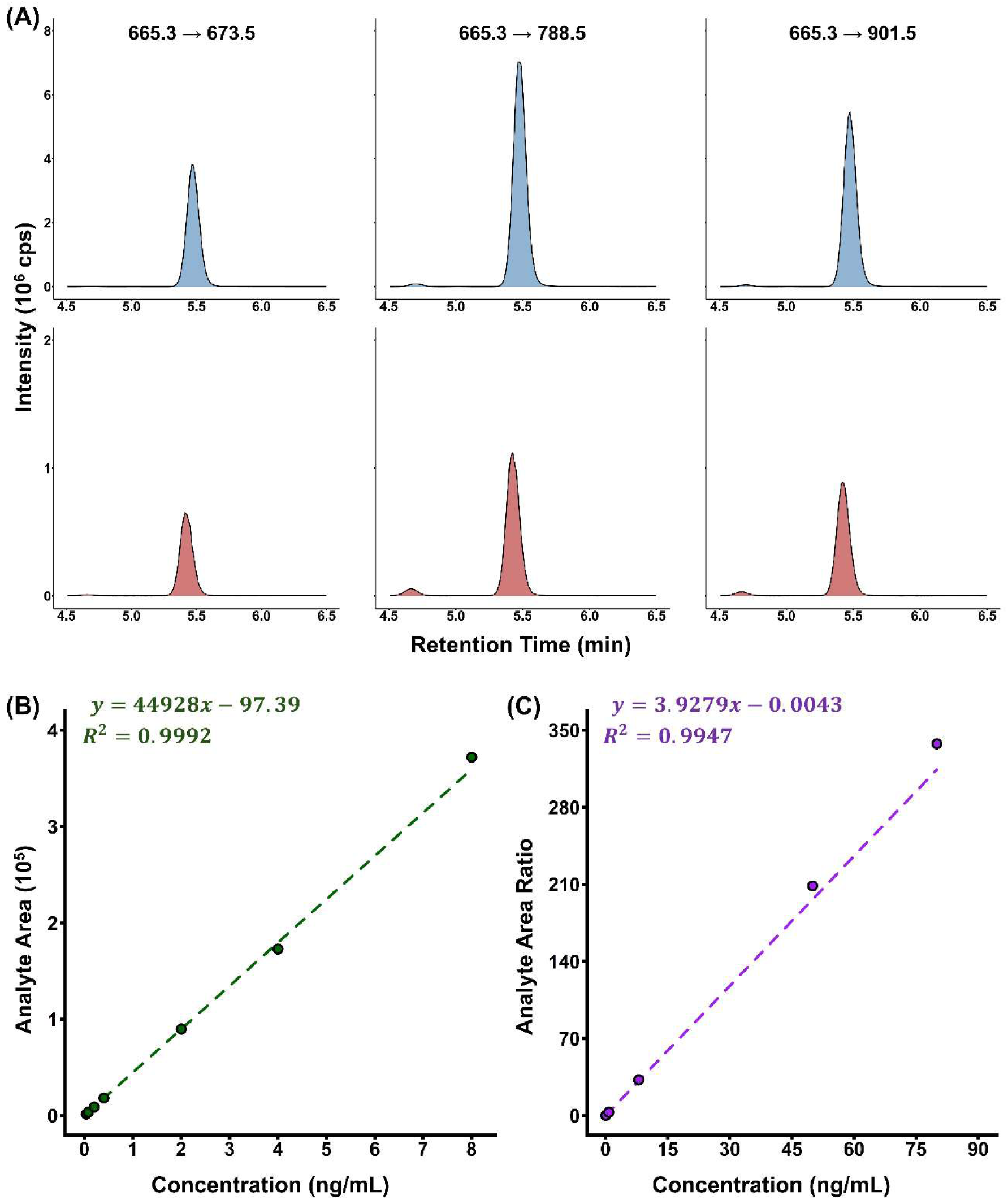
Multiple reaction monitoring confirmation and calibration curves for LUZ19 signature peptide EVAELDGQELAR. (A) Extracted MRM chromatograms for three monitored transitions in the light extended peptide calibrator at 80 ng/mL (blue) and a digested LUZ19 phage lysate at ∼10^9^ PFU/mL (red), confirming retention time and ion ratios between standard and sample. (B) Calibration curve for the synthetic (undigested) EVAELDGQELAR peptide across 0.04-8 ng/mL (single injection per level). (C) Calibration curve for the light extended peptide analog EVAELDGQELARKFD across 0.004-80 ng/mL, used to quantify digested phage samples during empirical copy number determination.

**Table 1:** Structural proteins and sequence coverage for LUZ19.

| Description | No. of Peptides | Coverage (%) | MW (kDa) |
|---|---|---|---|
| YP_001671977.1<br>major capsid protein | 15 | 55 | 37.6 |
| YP_001671995.1<br>hypothetical protein | 2 | 46 | 7.8 |
| YP_001671994.1<br>minor capsid protein | 1 | 25 | 8.7 |
| YP_001671965.1<br>hypothetical protein | 1 | 2 | 36.8 |

Peptide identification and selection were performed in E215, a taxonomically distinct myovirus relative to LUZ19. High-resolution mass spectrometry identified numerous E215 structural and hypothetical proteins, from which candidates were also filtered by structural annotation and abundance to four proteins for further evaluation (**Table S5**). Although the hypothetical protein candidate showed higher sequence coverage (59% vs. 21%), it was excluded from further consideration due to its unconfirmed functional annotation. The major capsid protein was selected instead as a target better suited to an initial feasibility assessment of assay translatability. Two candidate signature peptides, IQSQDQIDPK (**Figure S2**) and FITLGVEK (**Figure S2 and S3**), were identified from the E215 major capsid protein (YP_009619448.1). Neither peptide occurred in the PAO1 proteome.

### Determination of the Empirical Copy Number

The variability of the DLA method was assessed across three independent LUZ19 stocks (A, B, and C) prepared at concentrations ranging from 10^5^ to 10^9^ PFU/mL. Considerable variability in PFU/mL estimates was observed across stocks and dilution levels, with the coefficient of variation (CV) ranging from 9.6 to 59.6% and log-log slopes of DLA titer against dilution level of 1.01 (95% CI: 0.95 to 1.07), 1.05 (95% CI: 1.02 to 1.09), and 0.99 (95% CI: 0.95 to 1.02) for stocks A, B, and C respectively (**Figure 4A**). Corresponding MS analysis of the same phage lysate stocks demonstrated a near-proportional relationship between peak area ratio and phage concentration across all three stocks, with weighted R^2^ values of 0.9453, 0.8496, and 0.9434 for stocks A, B, and C respectively and log-log slopes of 1.02 (95% CI: 0.92 to 1.12), 0.89 (95% CI: 0.59 to 1.20), and 1.11 (95% CI: 1.03 to 1.20) (**Figure 4B**).

**Figure 4:**
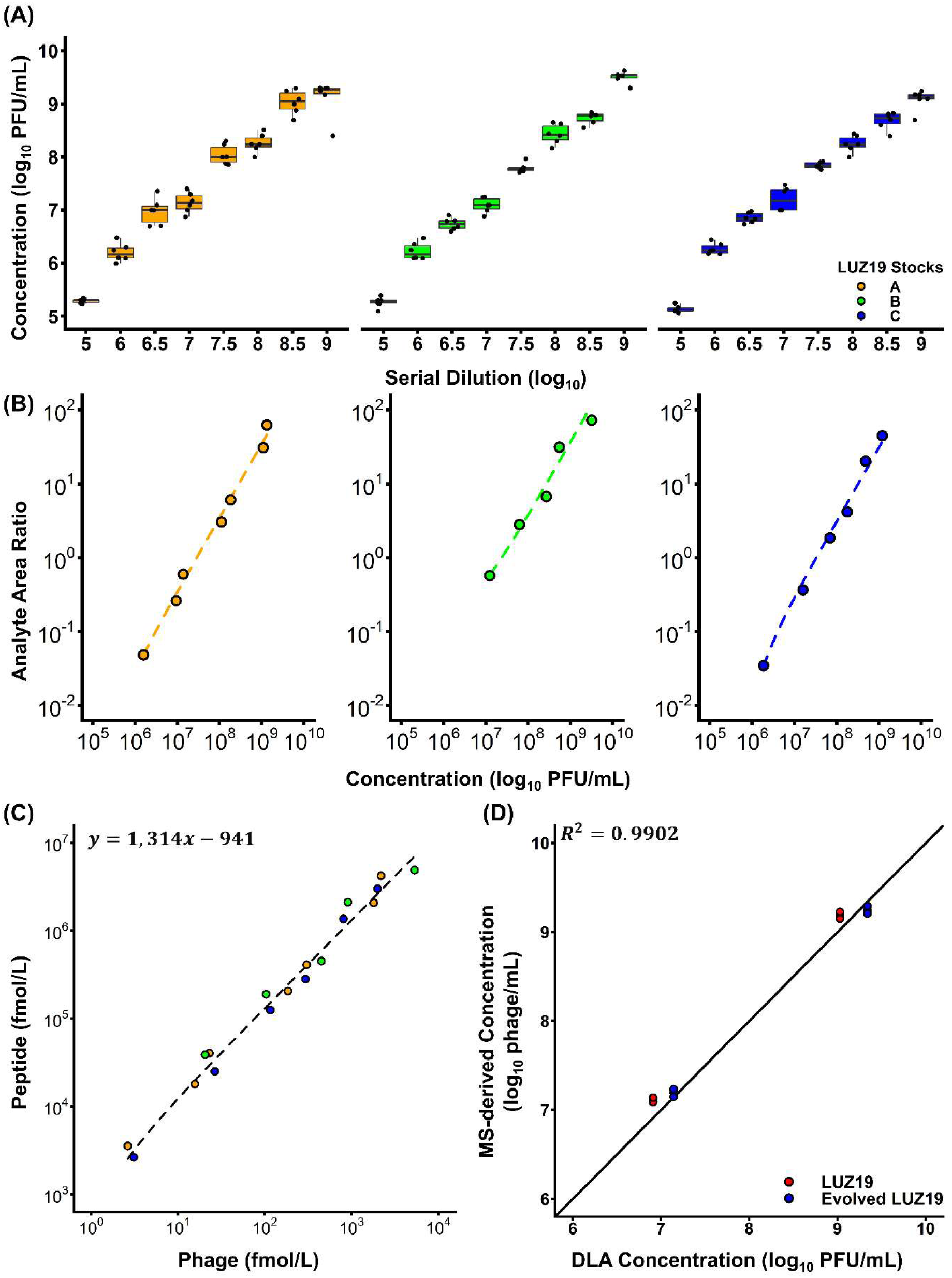
Quantitation of LUZ19 phage stocks by tandem mass spectrometry and comparison with the double layer agar assay. (A) Variability of DLA-derived PFU/mL estimates across three independent LUZ19 stocks (A, B, and C) prepared at 10-fold and half-log dilutions spanning 10^5^ to 10^9^ PFU/mL. Box plots represent the distribution of sextuplicate measurements at each dilution level, with individual observations overlaid. (B) Corresponding LC-MS/MS analysis of the same phage lysate stocks A, B, and C on a log-log scale where dashed lines represent weighted linear regression fits to the untransformed data for each stock. Lysates excluded from the regression (n = 6) are not shown. (C) Weighted Deming regression fitted to untransformed concentrations and displayed on logarithmic axes relating peptide molarity (fmol/L) derived from mass spectrometry to DLA-derived particle molarity (fmol/L) derived from the double layer agar assay across all three stocks, with titers converted to molarity using Avogadro’s number and peptide concentrations using the molecular weight of the signature peptide, yielding an empirical copy number of 1,314 (95% CI: 1,098 to 1,529) from 18 included lysates. (D) Agreement between mass spectrometry- and double layer agar assay-derived concentration estimates for unmodified LUZ19 (red) and evolved LUZ19 (blue) samples prepared at approximately 10^7^ and 10^9^ PFU/mL (10^5^ PFU/mL was below the LLOQ and excluded) and assessed in triplicate across three separate days. The solid black line is the line of identity. R^2^ is the correlation between mean log_10_ predicted and observed concentrations. Agreement was quantified as the mean log_10_ difference of 0.18 (1.52-fold [95% CI: 1.17 to 1.99]) for unmodified and −0.02 (0.95-fold [95% CI: 0.82 to 1.11]) for evolved LUZ19.

Peptide and phage molar concentrations were derived from mass spectrometry measurements and from plaque assay counts, respectively, and expressed in femtomolar units for regression analysis. An error ratio (λ = 0.076) was estimated from replicate QC injections and plaque titer replicates, reflecting superior precision of the mass spectrometry relative to the DLA assay. Deming regression of the peptide molarity against the DLA-derived particle molarity yielded an empirical copy number of 1,314 (95% CI: 1,098 to 1,529; n = 18 lysates), with stock specific estimates of 1,346, 1,257, and 1,181 for stocks A, B, and C (**Table S3**). The intercept was −941 fmol/L (95% CI: −3,238 to 1,356) and not distinguishable from zero, supporting a proportional relationship between peptide and particle molarity across the concentration range evaluated (**Figure 4C**). Levels below the LLOQ (∼ 10^5^ PFU/mL) and lysates outside the dilution-linearity criterion were excluded from the regression and the sensitivity of the empirical copy number to the inclusion criteria is shown in **Table S3**. For E215, the empirical copy number was 247 (95% CI: 210 to 284), determined from 10^8^ – 10^9^ PFU/mL (n = 5 lysates) using the FITLGVEK peptide (**Table S3 and Figure S3**). Applied to LUZ19 and E215 mixtures, measured concentrations were on average 2.6-fold (range 1.3 to 5.1) higher than expected values and 3.3-fold (range 1.3 to 8.6) higher than DLA titers determined on phage-resistant PAO1 mutants (**Figure S4 and Table S6**). No significant cross-phage interference was observed (LUZ19, β = 0.008, 95% CI: −0.006 to 0.022, p = 0.23; E215, β = −0.009, 95% CI: −0.091 to 0.072, p = 0.82).

### Evaluation of Quantitative Performance

Method performance was evaluated using three independent analytical runs prepared on separate days. Back-calculated concentrations of calibration standards met acceptance criteria across the calibration range of 0.004 to 80 ng/mL, with bias spanning −7.0 to 4.6% (**Table 2**). The light extended peptide QC samples at 0.006, 0.4, and 40 ng/mL demonstrated intra-day precision ranging from 0.5% to 9.3% CV (**Table 3**) and inter-day precision ranging from 6.6% to 12.0% CV (**Table 3**), with back-calculated accuracy spanning −19.5% to 6.1% bias across each concentration level and run. Pooled across runs, 78% of QC samples and at least 67% of replicates at each concentration level were within the ± 15% acceptance threshold, meeting this ICH M10 acceptance criterion. Chromatograms of blank culture medium and uninfected PAO1 culture lysate (host proteome blank) showed no interfering response at the retention time of the LUZ19 signature peptide (**Figure S5**).

**Table 2:**
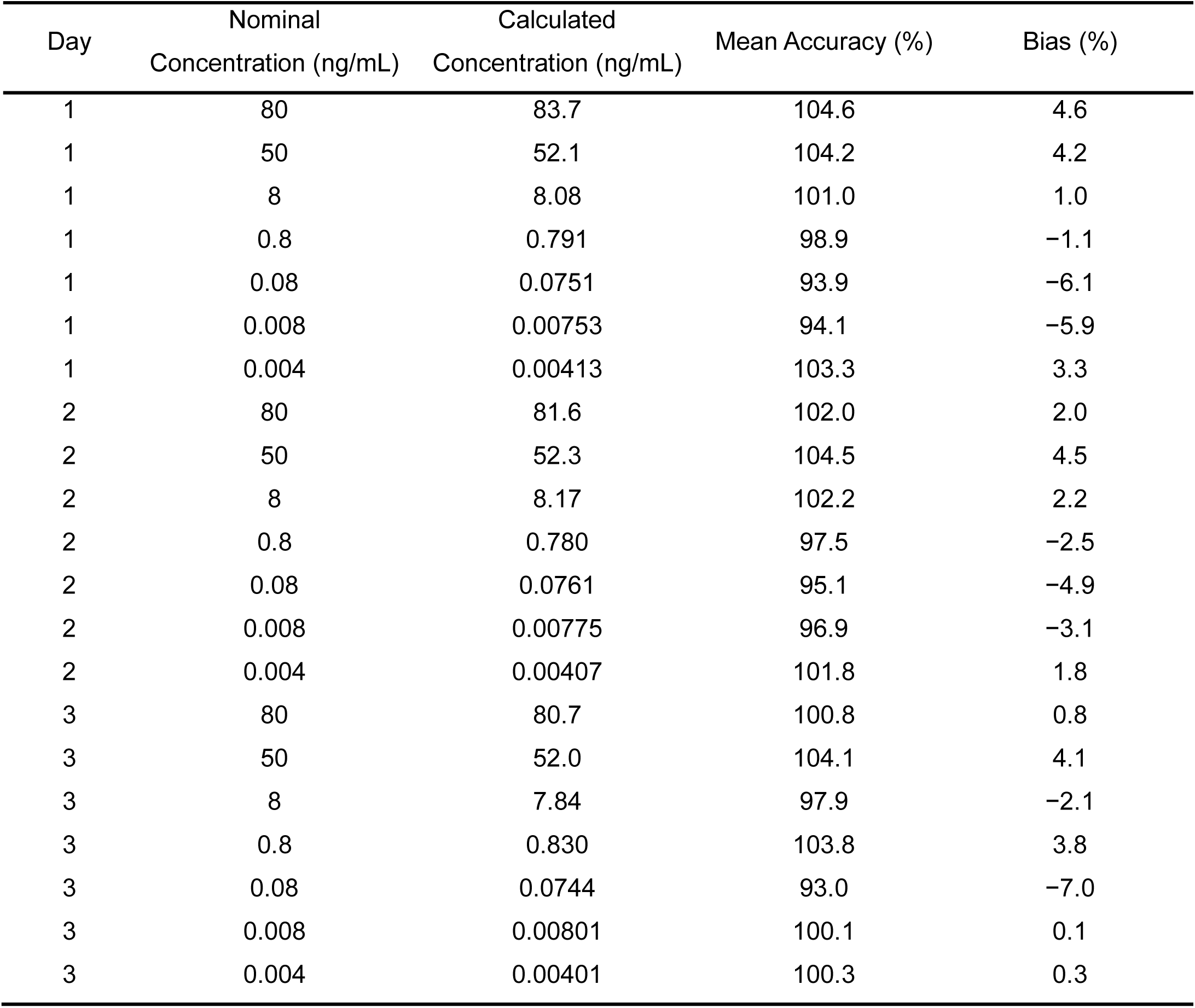
Back-calculated accuracy of calibration standards across three independent analytical runs.

| Day | Nominal<br>Concentration (ng/mL) | Calculated<br>Concentration (ng/mL) | Mean Accuracy (%) | Bias (%) |
|---|---|---|---|---|
| 1 | 80 | 83.7 | 104.6 | 4.6 |
| 1 | 50 | 52.1 | 104.2 | 4.2 |
| 1 | 8 | 8.08 | 101.0 | 1.0 |
| 1 | 0.8 | 0.791 | 98.9 | −1.1 |
| 1 | 0.08 | 0.0751 | 93.9 | −6.1 |
| 1 | 0.008 | 0.00753 | 94.1 | −5.9 |
| 1 | 0.004 | 0.00413 | 103.3 | 3.3 |
| 2 | 80 | 81.6 | 102.0 | 2.0 |
| 2 | 50 | 52.3 | 104.5 | 4.5 |
| 2 | 8 | 8.17 | 102.2 | 2.2 |
| 2 | 0.8 | 0.780 | 97.5 | −2.5 |
| 2 | 0.08 | 0.0761 | 95.1 | −4.9 |
| 2 | 0.008 | 0.00775 | 96.9 | −3.1 |
| 2 | 0.004 | 0.00407 | 101.8 | 1.8 |
| 3 | 80 | 80.7 | 100.8 | 0.8 |
| 3 | 50 | 52.0 | 104.1 | 4.1 |
| 3 | 8 | 7.84 | 97.9 | −2.1 |
| 3 | 0.8 | 0.830 | 103.8 | 3.8 |
| 3 | 0.08 | 0.0744 | 93.0 | −7.0 |
| 3 | 0.008 | 0.00801 | 100.1 | 0.1 |
| 3 | 0.004 | 0.00401 | 100.3 | 0.3 |

**Table 3:** Assessment of precision and accuracy.

| **Intra-day precision and accuracy.** |  |  |  |  |  |  |  |
|---|---|---|---|---|---|---|---|
| Day | QC Level | Nominal Concentration (ng/mL) | n | Mean Calculated Concentration (ng/mL) | CV (%) | Mean Accuracy (%) | Bias (%) |
| 1 | H | 40 | 3 | 38.9 | 1.3 | 97.2 | −2.8 |
| 1 | M | 0.4 | 3 | 0.349 | 1.6 | 87.2 | −12.8 |
| 1 | L | 0.006 | 3 | 0.00499 | 3.1 | 83.1 | −16.9 |
| 2 | H | 40 | 3 | 35.4 | 1.9 | 88.5 | −11.5 |
| 2 | M | 0.4 | 3 | 0.322 | 0.5 | 80.5 | −19.5 |
| 2 | L | 0.006 | 3 | 0.00539 | 9.3 | 89.9 | −10.1 |
| 3 | H | 40 | 3 | 42.1 | 0.7 | 105.2 | 5.2 |
| 3 | M | 0.4 | 3 | 0.374 | 0.6 | 93.6 | −6.4 |
| 3 | L | 0.006 | 3 | 0.00637 | 2.4 | 106.1 | 6.1 |
| **Inter-day precision and accuracy** |  |  |  |  |  |  |  |
| QC Level | Nominal Concentration (ng/mL) | n | Mean Calculated Concentration (ng/mL) | CV (%) | Mean Accuracy (%) | Bias (%) |  |
| H | 40 | 9 | 38.8 | 7.6 | 97.0 | −3.0 |  |
| M | 0.4 | 9 | 0.348 | 6.6 | 87.1 | −12.9 |  |
| L | 0.006 | 9 | 0.00558 | 12.0 | 93.1 | −6.9 |  |

To evaluate the accuracy of the empirical copy number, phage lysate QC samples from both the unmodified and evolved LUZ19 were prepared at approximately 10^5^, 10^7^, and 10^9^ PFU/mL and analyzed in triplicate across three separate days. Phage-equivalent concentrations estimated by the mass spectrometry method using the empirical copy number were compared with DLA-derived titers. At levels at or above the LLOQ, the mean difference was 0.18 log_10_ units (1.52-fold [95% CI: 1.17 to 1.99]) for unmodified LUZ19 and −0.02 log_10_ units (0.95-fold [95% CI: 0.82 to 1.11]) for evolved LUZ19 (**Figure 4D**). Lysates at ∼10^5^ PFU/mL were below the LLOQ and are not included in these statistics. At this level the peptide was not detected in 5 of 18 injections, and the mean of the detected values was below the LLOQ. EVAELDGQELAR was also identified in the evolved LUZ19 lysate by high resolution MS/MS (precursor mass error within 2 ppm), indicating retention of the target peptide in the evolved lineage (**Figure S6**). Predicted phage-equivalent concentrations had intra-day geometric CVs of 1.0 to 4.3% and inter-day geometric CVs of 5.5 to 9.0% across lysate levels. Bland-Altman analysis of LUZ19 QC lysate data at or above the LLOQ demonstrated mean differences of 0.18 (unmodified) and −0.02 (evolved) log_10_ units between MS-derived phage-equivalent and DLA-derived concentrations, with 95% limits of agreement of 0.03 (95% CI: −0.12 to 0.18) to 0.34 (95% CI: 0.19 to 0.49) log_10_ units for unmodified LUZ19 and −0.13 (95% CI: −0.23 to −0.02) to 0.09 (95% CI: −0.02 to 0.19) log_10_ units for evolved LUZ19 (**Figure S7**). Pooled across both lysates, the mean difference was 0.081 log_10_ (95% CI: 0.019 to 0.144), with limits of agreement of −0.052 (95% CI: −0.128 to 0.024) to 0.214 (95% CI: 0.138 to 0.290) log_10_ (**Figure S7**).

### Phage Directed Evolution and Host Range Expansion

Nine clinical *Pseudomonas aeruginosa* isolates obtained from the CDC/FDA AR isolate bank were used for directed evolution and efficiency of plating assessment. At Day 0, LUZ19 demonstrated variable EOP across the clinical isolate panel relative to the reference strain PAO1, with several isolates showing limited or no susceptibility (**Table S7**). Following directed evolution, EOP increased by at least one class for one isolate (AR-0254). AR-0257 increased approximately 390-fold but remained in the low EOP class (**Table S7**). The evolved LUZ19 demonstrated detectable lytic activity against two isolates that were resistant or poorly susceptible to the unmodified phage at Day 0.

## Discussion

The development of phage therapy as a clinically viable treatment modality has long been constrained by the absence of analytical methods capable of meeting the quantitative standards expected in modern drug development. As a proof-of-concept for a single therapeutic phage, this work demonstrates that targeted tandem mass spectrometry can provide reproducible, host-independent quantitation through detection of a structural signature peptide, anchoring phage enumeration within the established paradigm of bioanalytical method development. Current approaches to phage quantitation each measure fundamentally distinct biological properties and carry inherent tradeoffs that reflect the respective analytical scope of each method. The DLA assay quantifies infectious particles capable of completing a lytic cycle under defined conditions, providing biologically meaningful information directly relevant to therapeutic efficacy, but is labor-intensive, highly variable, and unsuitable for high-throughput workflows. qPCR measures genome copy number with high sensitivity and throughput, capturing a distinct and complementary biological property, though it does not differentiate between infectious and non-infectious particles. This inability to differentiate infectious from non-infectious particles can be particularly consequential for exposure-response characterization in PK/PD analyses where functional phage concentration governs bactericidal activity. LC-MS/MS occupies a complementary and distinct position in this landscape, providing a precise, reproducible measure of structural protein concentration that addresses the analytical precision and rigor that plaque-based and nucleic acid-based methods are not designed to provide. However, the DLA assay remains indispensable as the reference standard for infectivity, and the combination of these approaches offers a more comprehensive characterization of therapeutic phage preparations than any single method alone. This combined approach opens a pathway toward the rigorous PK/PD characterization needed for clinical translation.

A conceptual strength of the signature peptide approach lies in the selection of a target derived from a high-abundance structural capsid protein with a defined copy number per virion. The fixed stoichiometry of capsid proteins provides an intrinsic signal that is directly proportional to phage concentration, offering a stable and structurally invariant quantitation anchor. Unlike tail fiber proteins, which may exhibit variable interactions with bacterial hosts and introduce additional sources of analytical noise, capsid proteins are not subject to the same host-dependent variability, making them well-suited targets for reproducible quantitation across diverse sample types. Furthermore, while PCR-based approaches require independent optimization of amplification conditions for each phage target, a requirement that scales poorly with cocktail complexity, LC-MS/MS enables simultaneous quantitation of multiple phages by selecting unique signature peptides with distinct MRM transitions. This represents a fundamentally different relationship between measured signal and biological target, one that is directly proportional to virion concentration and uniquely suited to the multiplexed quantitation demands of an arbitrary N-phage cocktail.

The practical implications of this approach extend directly to the clinical scenarios where current analytical methods have been unable to fully meet the demands of phage PK/PD characterization. The ELIMINATE trial, which evaluated a six-phage cocktail, illustrates this challenge where quantitation required three high-efficiency host strains and yielded only a composite titer. While a pragmatic solution given available methods, it does not support the individual phage PK/PD characterization needed for cocktail optimization.^29^ LC-MS/MS would address this gap by enabling host-independent, simultaneous quantitation of each phage component within a single analytical run. Unlike conventional small molecule drugs, phages must reach the site of infection to interact with susceptible bacteria, meaning that plasma concentrations alone are insufficient to characterize therapeutic exposure. Quantitation at the site of infection, whether in pulmonary tissue, wound fluid, or other relevant compartments, is essential for understanding the exposure-response relationships that govern bactericidal activity.^30^ The matrix flexibility and analytical specificity of LC-MS/MS make it particularly well-suited for tissue-level quantitation, where host-dependent methods face compounding sources of variability from endogenous biological components, further limiting already variable quantitative performance.

A significant consideration in evaluating the agreement between LC-MS/MS and DLA-derived concentration estimates is the inherent analytical floor of the plaque assay itself. Plaque count data follows a Poisson distribution, which introduces an irreducible baseline variability that is independent of operator skill or procedural standardization. This Poisson-derived variability establishes a theoretical minimum CV for the DLA that cannot be overcome regardless of how rigorously the assay is performed and was reflected in the CV values observed across stocks in this study. Anderson and colleagues proposed a 0.33 log equivalence threshold as the criterion for agreement between replicates of the DLA, acknowledging that differences within this range are indistinguishable from the natural variability of plaque-based enumeration.^9^ A difference of 0.33 log_10_ units corresponds to a 2.14-fold difference in concentration. For both unmodified and evolved LUZ19, the bias and its 95% confidence interval fell within this threshold, and the limits of agreement were within it for evolved LUZ19 and near it for unmodified LUZ19, indicating agreement with the plaque assay within the variability the plaque assay itself exhibits. This inherent variability floor also has direct implications for experimental design. The imprecision of plaque-based enumeration necessitates replicate measurements to achieve reliable concentration estimates, whereas the analytical precision of mass spectrometry is sufficient to support singlet analysis, making it amenable to high-throughput workflows.

The empirical copy number links peptide molarity to DLA-derived particle concentration. It reflects the number of copies of the target protein per virion, the ratio of total to infectious particles in the preparation, and the combined efficiency of capsid disruption, digestion, and peptide recovery. These terms may differ between phages and preparations, so the empirical copy number would scale with the structural copy number, serving as a proxy for it. Validation against phages with published capsid stoichiometry would be required before the empirical copy number is interpreted structurally. The consistency of stock-specific estimates across three independent phage stocks, combined with an intercept that was indistinguishable from zero, supports the proportional relationship between peptide and particle molarity across the evaluated concentration range. It should be noted, however, that PFU-based quantitation selectively enumerates infectious particles capable of completing a lytic cycle on the host isolate and may underestimate total virion concentration if non-infectious or defective particles are present in the lysate preparation. The concordance observed for evolved LUZ19 is a particularly meaningful finding in the context of phage therapy development, where directed evolution has been explored to generate phages with expanded therapeutic host ranges. Following directed evolution using a single-phage Appelmans protocol, LUZ19 demonstrated increased EOP against two clinical isolates that were resistant or poorly susceptible to the parental phage. The narrow host range of individual phages has historically limited the utility of phage therapy, driving the development of strategies such as phage cocktails, engineering of receptor binding proteins, and directed evolution to broaden clinical applicability.^31, 32^ A fundamental advantage of phage therapy over conventional antibiotics lies in this adaptability, whereas small molecule antibiotics are static entities that cannot adapt as resistance emerges. Phages can be iteratively evolved to overcome emerging resistant strains, offering a renewable therapeutic modality. In a single evolved LUZ19 lineage, the target peptide was retained and quantified, with agreement to DLA similar to the parental phage after 10 rounds of selection. This supports the broader applicability of the assay to phage variants and suggests that capsid proteins may represent stable quantitation anchors even under selective evolutionary pressure. This approach was also applied in a second, taxonomically distant phage, E215, for which discovery-phase identification yielded two candidate signature peptides. An empirical copy number was determined for FITLGVEK and used to quantify E215 in mixed samples with LUZ19, supporting the feasibility of extending this strategy to a second phage and concurrent quantitation.

Several limitations of this proof-of-concept study warrant explicit acknowledgement. This work was conducted using a single therapeutic phage, LUZ19, and a single signature peptide derived from the major capsid protein. Peptide identification, detectability, and the empirical copy number were determined in a second phage and applied in mixed samples, but accuracy and precision of E215 measurements were not assessed. The multiplexed quantitation of LUZ19 and E215 was evaluated only as a preliminary feasibility assessment, and further optimization is required before quantitation in mixtures can be considered accurate. Generalizability to other phage morphologies and structural proteins beyond these two targets remains an important avenue for future investigation. The empirical copy number is derived from a proportional relationship between detectable peptide molarity and DLA-derived particle concentration and is an analytical scaling factor rather than a structurally verified copy number. Its interpretation as a structural copy number would require validation against phages with published capsid stoichiometry or orthogonal structural methods such as cryo-electron microscopy or native mass spectrometry. The extended-peptide internal standard is added before digestion and corrects for tryptic digestion and instrument variability, but not for capsid disruption, precipitation recovery, or cleavage-site accessibility in the denatured protein, which were not independently measured. However, SIL full-length phage proteins or virions that would capture this variability are not currently available and remain a limitation in the field. The phage-equivalent LLOQ is derived from the lowest peptide calibrator and may not reflect the true LLOQ of intact phage lysates. Because the empirical copy number is calibrated to DLA and agreement was assessed against DLA, the comparison demonstrates consistency with the reference method and not independent measurement of absolute phage concentration. Additionally, the current assay was developed and evaluated in phage lysate and has not yet been tested in complex biological matrices relevant to clinical practice, such as plasma, urine, or tissue homogenates. Matrix-specific interferences in these sample types may affect assay performance and will require dedicated method development prior to clinical implementation. Finally, because the phage-equivalent concentration is calibrated to PFU/mL, it may differ from total virion concentration if non-infectious or defective particles are present in significant proportions, which an orthogonal total-particle method would be needed to resolve.

Looking forward, several directions present themselves as natural extensions of this work. A logical and immediate next step is the transition toward a fully functional tandem mass spectrometry assay, in which multiple signature peptides derived from the same structural protein, or from multiple structural proteins simultaneously, are monitored within a single analytical run. Such an approach would provide not only redundant quantitative targets that strengthen confidence in concentration estimates, but also a comprehensive analytical fingerprint of the virion that confirms structural integrity, composition, and phage identity in parallel. Agreement between independent peptide-derived concentration estimates would provide strong orthogonal evidence for the reliability of the empirical copy number, and internal consistency checks across multiple targets would substantially improve the robustness and clinical readiness of the assay. Translation of the assay to complex biological matrices encountered in clinical practice represents another important next step, as quantitation in clinically relevant compartments would support not only pharmacokinetic characterization of phage distribution but also pharmacodynamic assessment of bactericidal activity at the site of infection. These environments introduce matrix-specific interferences that will require characterization and appropriate sample preparation strategies such as solid-phase extraction. The lowest LUZ19 level included in the quantitative analyses was approximately 10^6^ PFU/mL using analytical flow, with a calibrator derived phage-equivalent LLOQ of 5.685 log_10_ phage/mL, which falls within a clinically relevant range for therapeutic phage concentrations expected during treatment.^29^ Should higher sensitivity be required for specific applications, transitioning to a nano-flow or micro-flow LC configuration would be expected to improve and extend the quantitation range by one to two orders of magnitude. Finally, perhaps the most transformative future direction enabled by this work is the extension to multiplexed enumeration of phage cocktails. By selecting phage-specific signature peptides from each component of a cocktail and monitoring unique MRM transitions simultaneously, individual phage concentrations could be resolved within a single analytical run. This multiplexing capability is fundamentally beyond the reach of existing enumeration methods and would be transformative for cocktail PK characterization and clinical monitoring.

## Conclusion

Together, these findings position mass spectrometry-based signature peptide quantitation as a promising complement to existing methods for phage enumeration. By anchoring quantitation to a structural protein within the framework of bioanalytical mass spectrometry, this approach offers a path toward the analytical standards expected in modern drug development. As phage therapy advances towards clinical application, continued validation and extension of this strategy may help support the pharmacokinetic and pharmacodynamic characterization that its development will require.

## Supporting information

Peptide reagents and LUZ19 MRM transitions, E215 singleplex and multiplex LC-MS/MS methods, discovery-phase identification, structural protein coverage, and targeted MRM analysis of E215 signature peptide, sensitivity of the empirical copy number to lysate inclusion and stock-specific estimates, LUZ19 ion ratios, back-calculated calibration bias, multiplexed quantification of LUZ19 and E215 in mixed samples, blank and host proteome chromatograms, the MS/MS spectrum of the evolved LUZ19 peptide, Bland-Altman analysis of predicted versus observed LUZ19 concentrations, and efficiency of plating after directed evolution.

## Notes

The authors declare no competing financial interest.

## Acknowledgements

This study is supported by the National Institute of Allergy and Infectious Diseases of the National Institutes of Health under award numbers R01AI177997 and L30AI194397.

We thank Dr. Alan Friedman (University at Buffalo, Department of Materials Design and Innovation) for instrumentation support related to the LTQ Orbitrap XL (Thermo Fisher Scientific, Waltham, MA, USA).

