## Supporting Information for "Translational Quantitative Proteomic Assay for Bacteriophages: Opening a Frontier in Phage Pharmaceutical Development"

**Table S1: Peptide reagents for LUZ19 and E215**

| Peptide | Sequence | Label | MW (Da) | Role |
|---|---|---|---|---|
| LUZ19 signature peptide | EVAELDGQELAR | - | 1329.43 | MS optimization; analyte |
| LUZ19 light extended peptide | EVAELDGQELARKFD | - | 1719.87 | Calibrator |
| LUZ19 SIL extended peptide | EVAELDGQELA(R)KFD | [$^{13}C_6$, $^{15}N_4$]-Arg, +10 Da | 1729.80 | Internal standard |
| E215 signature peptide | FITLGVEK | - | 906.09 | MS optimization; analyte |
| E215 light extended peptide | QSKFITLGVEKRAK | - | 1604.91 | Calibrator |
| E215 SIL extended peptide | QSKFITLGVE(K)RAK | [$^{13}C_6$, $^{15}N_2$]-Lys, +8 Da | 1612.85 | Internal standard |

*Residues in parentheses carry the stable isotope label. MW: calculated average molecular weight. SIL: stable isotope-labeled.*

**Table S2: LUZ19 MRM transitions.**

| Compound | Q1 (*m/z*) | Q3 (*m/z*) | Ion | Role | CE (V) | Dwell (ms) | Expected ion ratio |
|---|---|---|---|---|---|---|---|
| EVAELDGQELAR | 665.3 | 788.5 | y7 | Quantifier | 32 | 150 | 1.00 |
| EVAELDGQELAR | 665.3 | 901.5 | y8 | Qualifier | 32 | 150 | 0.74 |
| EVAELDGQELAR | 665.3 | 673.5 | y6 | Monitored | 32 | 150 | - |
| SIL internal standard | 670.5 | 798.2 | y7 | Quantifier | 32 | 150 | - |
| SIL internal standard | 670.5 | 1040.4 | y9 | Qualifier | 32 | 150 | - |

*y7 was selected as the quantifier as the most intense transition. Expected ratio is the mean calibrator ratio.*

### Supplementary Methods

*LC-MRM-MS Analysis*

E215 peptides were separated on a Waters Premier CSH C18 column (2.1 mm × 100 mm, 3.5 µm, Waters, Milford, MA, USA). A 15-minute gradient was employed with a flow rate of 0.200 mL/min. Injection volumes ranged from 5 to 25 µL, with larger volumes used at lower concentrations. Mobile phases consisted of 10% methanol in water containing 0.1% formic acid (Mobile Phase A) and 95% acetonitrile in water containing 0.1% formic acid (Mobile Phase B). The gradient was initiated at 5% B and held until 0.5 min, increasing to 15% B at 3.5 min, 20% B at 4.5 min, 35% B at 5 min, and 50% B at 5.5 min, followed by a step to 95% B at 6 min held until 10 min, and re-equilibrated at 5% B from 10.1 to 15 min. The column temperature was 40°C and the autosampler temperature was 8°C. The spray voltage was 2.5 kV, and the source temperature was 415°C. Quantitation of E215 was performed using multiple reaction monitoring of the transitions *m/z* 453.8 → 646.4 [$(M+2H)^{2+}$ → $y_6$, quantifier], and *m/z* 453.8 → 233.0 [$(M+2H)^{2+}$ → $a_2$, qualifier] for the signature peptide, and *m/z* 457.9 → 654.3 [$(M+2H)^{2+}$ → $y_6$, quantifier], and *m/z* 457.9 → 232.4 [$(M+2H)^{2+}$ → $a_2$, qualifier] for the SIL internal standard. For E215, the optimized collision energy was set to 20 V, the Q0 dissociation was set to 25 V (analyte) and 20 V (IS), and dwell time was 150 ms for each transition.

Mixed LUZ19 and E215 peptides were separated on a Waters Premier CSH C18 column (2.1 mm × 100 mm, 3.5 µm, Waters, Milford, MA, USA). A 15-minute gradient was employed with a flow rate of 0.230 mL/min. Injection volumes ranged from 5 to 25 µL, with larger volumes used at lower concentrations. The gradient was initiated at 9% B and held until 1 min, increasing to 10% B at 3.5 min (held until 4.5 min), 35% B at 5 min (held until 5.5 min), and 50% B at 6 min, followed by a step to 95% B at 6.1 min held until 10 min, and re-equilibrated at 9% B from 10.1 to 15 min. Both phages were monitored in a single method, using the transitions and mass spectrometer parameters of the single-plex assays.

Calibration standards contained both phages, each at its own concentration level, with each SIL internal standard at 1 ng/mL. To evaluate quantitation in mixed samples, LUZ19 and E215 lysates were each diluted 1:10, titered by DLA, and combined in equal volumes at prespecified concentrations to generate a 3 × 3 concentration grid. Expected titers were calculated as half the DLA titer of each phage. For each mixture, 285 µL was processed, reconstituted in 100 µL, and analyzed in triplicate. To independently quantify each phage in the mixtures, mixtures were also titered by DLA on phage-resistant PAO1 mutants. Calibration curves were fit by weighted linear regression ($1/x^2$), and peptide concentrations were converted to phage/mL using the empirical copy number determined for each phage. Phage-equivalent concentrations were compared with expected and resistant-mutant DLA titers as $\log_{10}$ differences, and cross-phage interference was assessed by regressing $\log_{10}$ phage-equivalent concentrations on the $\log_{10}$ expected titer of the same phage with the $\log_{10}$ expected titer of the other phage as an additional term.

**Table S3: Sensitivity of empirical copy number to lysate inclusion.**

| Analysis | Lysates (n) | Copy Number | Intercept (fmol/L) |
|---|---|---|---|
| **LUZ19** | | | |
| All lysates | 24 | 1,245 (1,070 – 1,420) | 428 (130 – 725) |
| Lysates ≥ LLOQ | 21 | 1,315 (1,107 – 1,523) | −818 (−2,300 – 664) |
| Primary analysis | 18 | 1,314 (1,098 – 1,529) | −941 (−3,238 – 1,356) |
| Stock A | 7 | 1,346 (1,002 – 1,690) | −78 (−4,374 – 4,219) |
| Stock B | 5 | 1,257 (243 – 2,271) | 14,064 (−88,030 – 116,159) |
| Stock C | 6 | 1,181 (728 – 1,635) | −1,106 (−22,947 – 20,735) |
| **E215** | | | |
| All lysates | 18 | 241 (174 – 308) | 19,836 (11,809 – 27,864) |
| Lysates ≥ LLOQ | 18 | 241 (174 – 308) | 19,836 (11,809 – 27,864) |
| Primary analysis | 5 | 247 (210 – 284) | 20,399 (−12,640 – 53,439) |

*Values are estimates (95% CI). Weighted Deming regression of peptide molarity on DLA-derived particle molarity (fmol/L). Primary analysis (dilution-linear lysates within stock) is the empirical copy number reported. LLOQ – lower limit of quantification.*

**Table S4: LUZ19 ion ratio for calibrators, peptide QCs, and lysates.** Mean ratio, standard deviation, and number of injections within ± 20% of the mean calibrator ratio for the qualifier-to-quantifier peak area ratio.

| Group | n | Mean ratio | SD | Within ± 20% |
|---|---|---|---|---|
| Calibrators | 7 | 0.743 | 0.030 | 7/7 |
| Peptide QCs | 9 | 0.776 | 0.037 | 9/9 |
| Unmodified lysate (~$10^7$ and $10^9$ PFU/mL) | 6 | 0.775 | 0.012 | 6/6 |
| Evolved lysate (~$10^7$ and $10^9$ PFU/mL) | 6 | 0.756 | 0.012 | 6/6 |

*Lysates at ~ $10^5$ PFU/mL were below the LLOQ and not included.*

**Figure S1: Back-calculated bias of calibration curves.** Percent bias of back-calculated concentrations from $1/x^2$ weighted calibration fits for (A) the LUZ19 synthetic peptide (0.04 – 8 ng/mL), (B) the LUZ19 light extended peptide calibrators (0.004 – 80 ng/mL), (C) the E215 synthetic peptide (0.02 – 4 ng/mL), and (D) the E215 light extended peptide calibrators (0.025 – 25 ng/mL). The solid line marks 0% bias and dashed lines ± 15%. Panels B and D are from the calibration run acquired with the lysates used for empirical copy number determination.

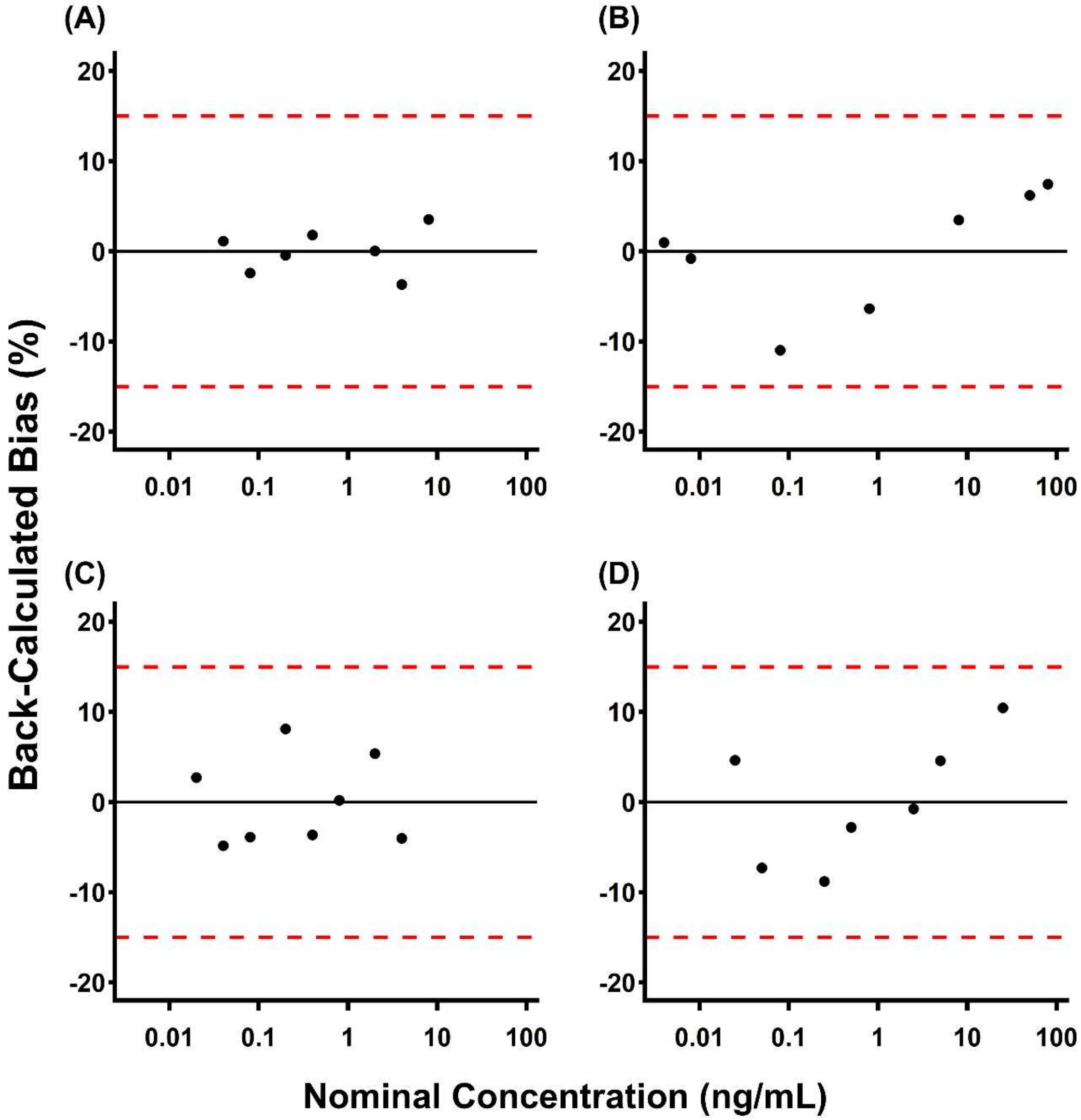

**Table S5: Structural proteins and sequence coverage for E215.**

| Description | No. of Peptides | Coverage (%) | MW (kDa) |
|---|---|---|---|
| YP_009619476.1 hypothetical protein | 16 | 59 | 32.2 |
| YP_009619448.1 major capsid protein | 5 | 21 | 41.7 |
| YP_009619454.1 tail sheath | 10 | 40 | 53.6 |
| YP_009619477.1 hypothetical protein | 6 | 39 | 23.4 |

**Figure S2: Discovery-phase identification of the E215 tryptic peptide IQSQDQIDPK by high-resolution Orbitrap mass spectrometry.** (A) Extracted ion chromatogram of the precursor ion corresponding to IQSQDQIDPK (m/z = 586.3013), showing a single chromatographic peak at retention time of ~22.3 min. (B) MS2 fragmentation spectrum of the precursor ion, with y-ion fragments (y2-y8) labeled in red, confirming peptide identity. (C) Sequence coverage map of the E215 major capsid protein, with identified tryptic peptides underlined. IQSQDQIDPK and FITLGVEK are boxed in red as candidate signature peptides for E215 quantitation.

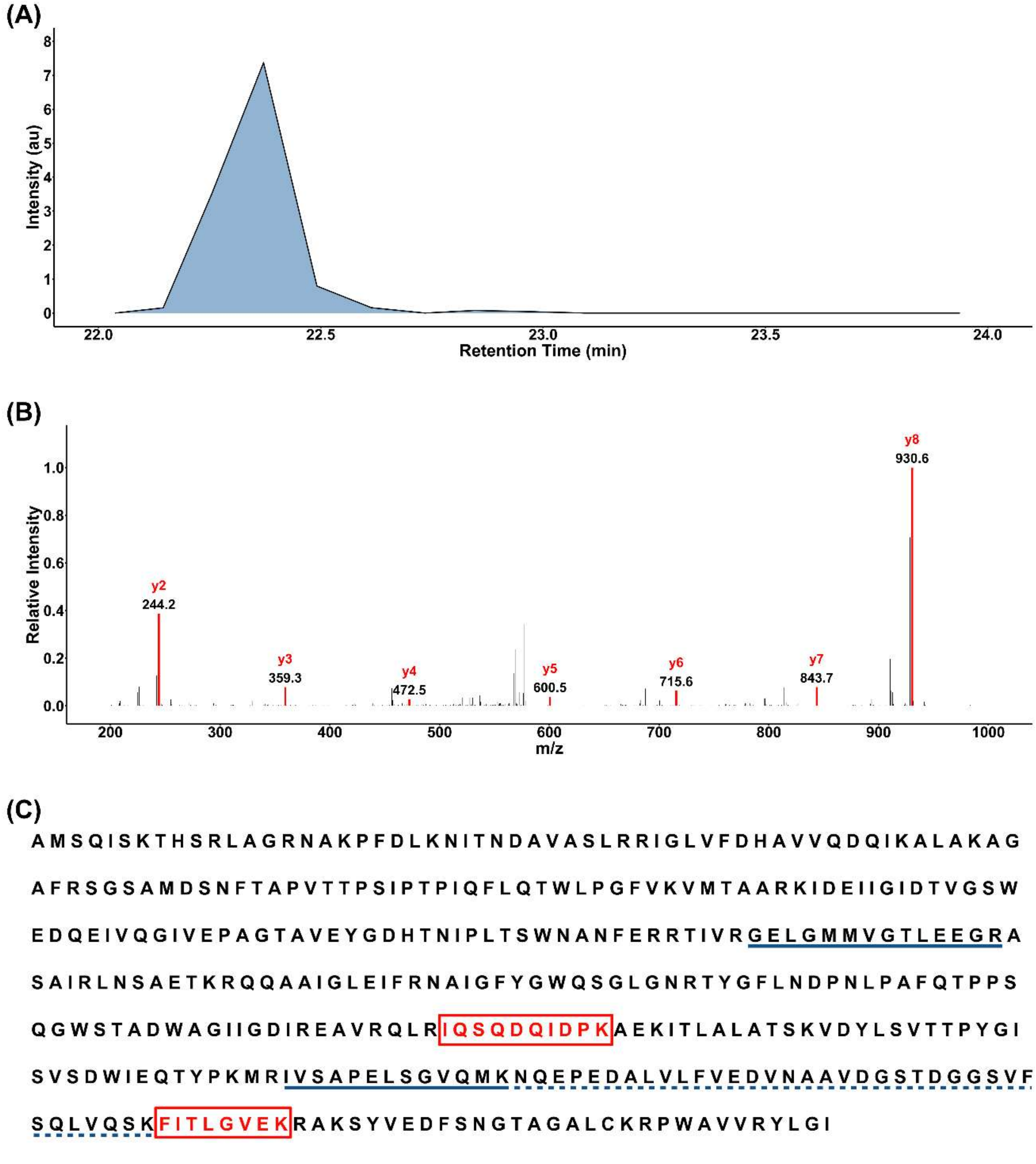

**Figure S3: Targeted MRM analysis of the E215 signature peptide.** (A) Representative MRM chromatograms for the E215 signature peptide FITLGVEK (precursor m/z 453.8) for the 25 ng/mL calibrator (blue) and an E215 lysate (red), showing the qualifier (453.8 → 233.0) and quantifier (453.8 → 646.4). (B) Synthetic peptide calibration curve for FITLGVEK (0.02 – 4 ng/mL) plotted as peak area (106) versus concentration. (C) Calibration curve from the empirical copy number analysis (0.025 – 25 ng/mL) plotted as the analyte/IS area ratio versus concentration.

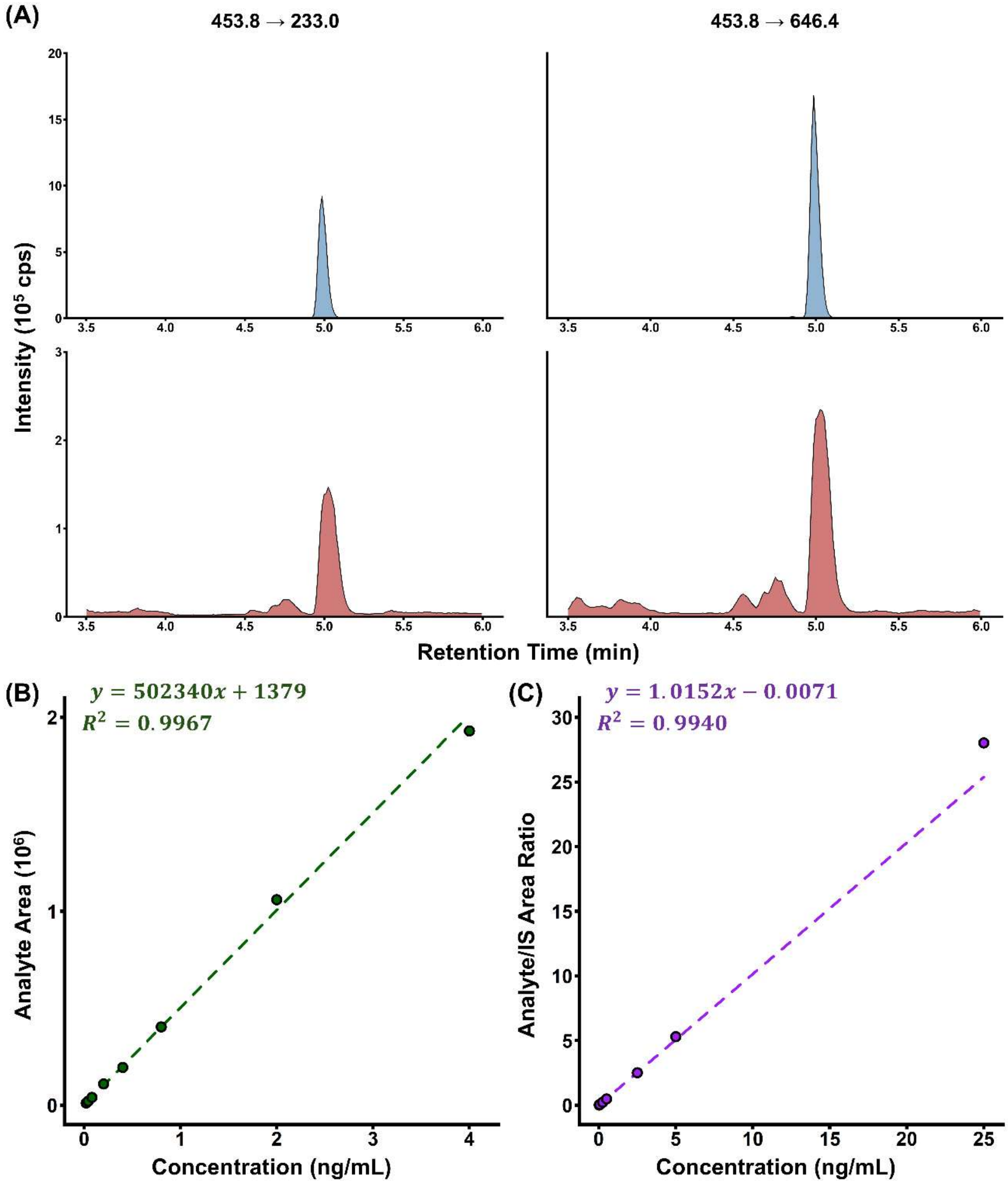

**Figure S4: Multiplexed LC-MRM quantification of LUZ19 and E215 in mixed samples.** (A) Measured versus expected titer for nine mixtures of LUZ19 and E215 (3 × 3 design) by MS (left) and by DLA on phage-resistant PAO1 mutants (right). Expected titers were derived from DLA of the single-phage stocks. Mean bias (measured – expected) was +0.41 for MS and −0.11 for DLA. The co-occurring phage did not significantly affect measured titer (LUZ19, $p = 0.23$; E215, $p = 0.82$). A sensitivity analysis using the nine mixture means gave $p = 0.47$ (LUZ19) and $p = 0.91$ (E215). (B) Representative MRM chromatograms of the high concentration (~$10^9$ PFU/mL) LUZ19 and E215 mixture showing quantifier (top) and qualifier (bottom) transitions for the LUZ19 peptide EVAELDGQELAR (left) and the E215 peptide FITLGVEK (right). Interference was tested by regressing each phage's $\log_{10}$ concentration on its own expected titer and the co-occurring phage's expected titer ($n = 27$ preparations per phage).

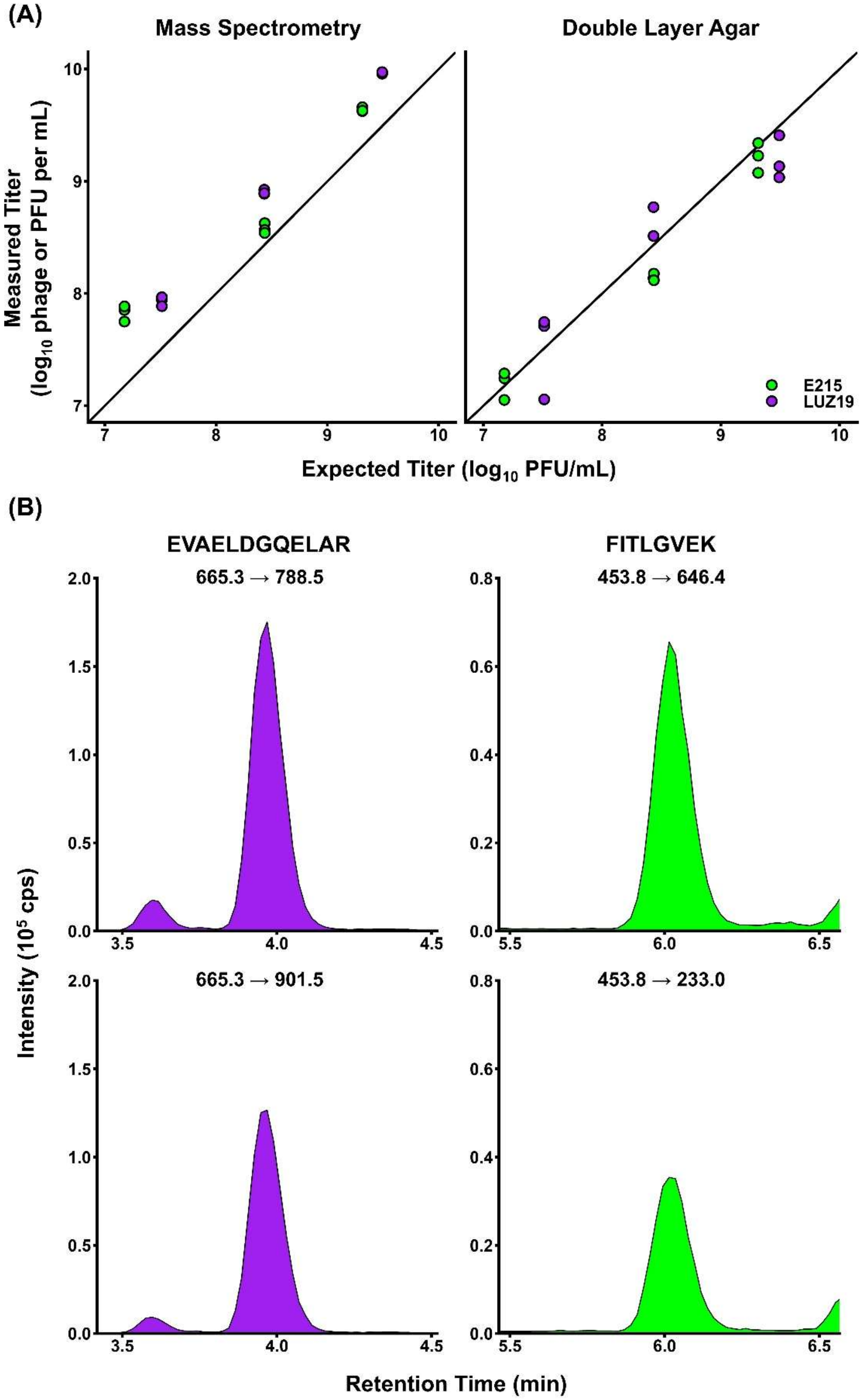
(A)
Mass Spectrometry
Double Layer Agar
Measured Titer
(log$_{10}$ phage or PFU per mL)
Expected Titer (log$_{10}$ PFU/mL)
E215
LUZ19
(B)
EVAELDGQELAR
FITLGVEK
665.3 → 788.5
453.8 → 646.4
665.3 → 901.5
453.8 → 233.0
Intensity (10$^5$ cps)
Retention Time (min)

**Table S6: Titers of LUZ19 and E215 in mixed samples by multiplexed LC-MRM and DLA.**

| Mixture | Expected | DLA (mean ± SD) | MS (mean ± SD) | MS − expected | DLA − expected |
|---|---|---|---|---|---|
| **LUZ19 (EVAELDGQELAR)** | | | | | |
| A1 | 9.49 | 9.03 ± 0.05 | 9.97 ± 0.01 | +0.47 | −0.46 |
| A2 | 8.43 | 8.51 ± 0.41 | 8.89 ± 0.01 | +0.46 | +0.08 |
| A3 | 7.51 | 7.71 ± 0.17 | 7.94 ± 0.03 | +0.43 | +0.20 |
| A4 | 9.49 | 9.13 ± 0.07 | 9.96 ± 0.00 | +0.47 | −0.36 |
| A5 | 8.43 | 8.14 ± 0.05 | 8.92 ± 0.01 | +0.49 | −0.30 |
| A6 | 7.51 | 7.06 ± 0.05 | 7.97 ± 0.01 | +0.45 | −0.46 |
| A7 | 9.49 | 9.41 ± 0.14 | 9.97 ± 0.02 | +0.48 | −0.08 |
| A8 | 8.43 | 8.77 ± 0.14 | 8.89 ± 0.02 | +0.46 | +0.34 |
| A9 | 7.51 | 7.75 ± 0.20 | 7.89 ± 0.04 | +0.37 | +0.23 |
| **Mean bias (SD)** | | | | +0.45 (0.03) | −0.09 (0.31) |
| **E215 (FITLGVEK)** | | | | | |
| A1 | 9.31 | 9.07 ± 0.09 | 9.62 ± 0.04 | +0.31 | −0.24 |
| A2 | 9.31 | 9.34 ± 0.18 | 9.66 ± 0.04 | +0.34 | +0.03 |
| A3 | 9.31 | 9.23 ± 0.18 | 9.63 ± 0.03 | +0.31 | −0.09 |
| A4 | 8.44 | 8.12 ± 0.31 | 8.63 ± 0.02 | +0.19 | −0.32 |
| A5 | 8.44 | 8.18 ± 0.08 | 8.57 ± 0.02 | +0.13 | −0.26 |
| A6 | 8.44 | 8.12 ± 0.08 | 8.54 ± 0.02 | +0.10 | −0.32 |
| A7 | 7.18 | 7.05 ± 0.13 | 7.75 ± 0.05 | +0.57 | −0.12 |
| A8 | 7.18 | 7.24 ± 0.08 | 7.85 ± 0.04 | +0.68 | +0.07 |
| A9 | 7.18 | 7.29 ± 0.09 | 7.88 ± 0.03 | +0.71 | +0.11 |
| **Mean bias (SD)** | | | | +0.37 (0.23) | −0.13 (0.17) |

*Values are $\log_{10}$ titers (measured – expected for bias). Expected and DLA values are PFU/mL; MS values are phage/mL. Expected: derived from DLA of the single-phage stocks. DLA: titer on phage-resistant PAO1 mutants.*

**Figure S5: Quantifier transition chromatograms of blank and host proteome blank samples.** Chromatograms of a blank (top) and a host proteome blank (bottom) at the retention times of the LUZ19 peptide EVAELDGQELAR (left) and the E215 peptide FITLGVEK (right). Red lines mark the expected retention times.

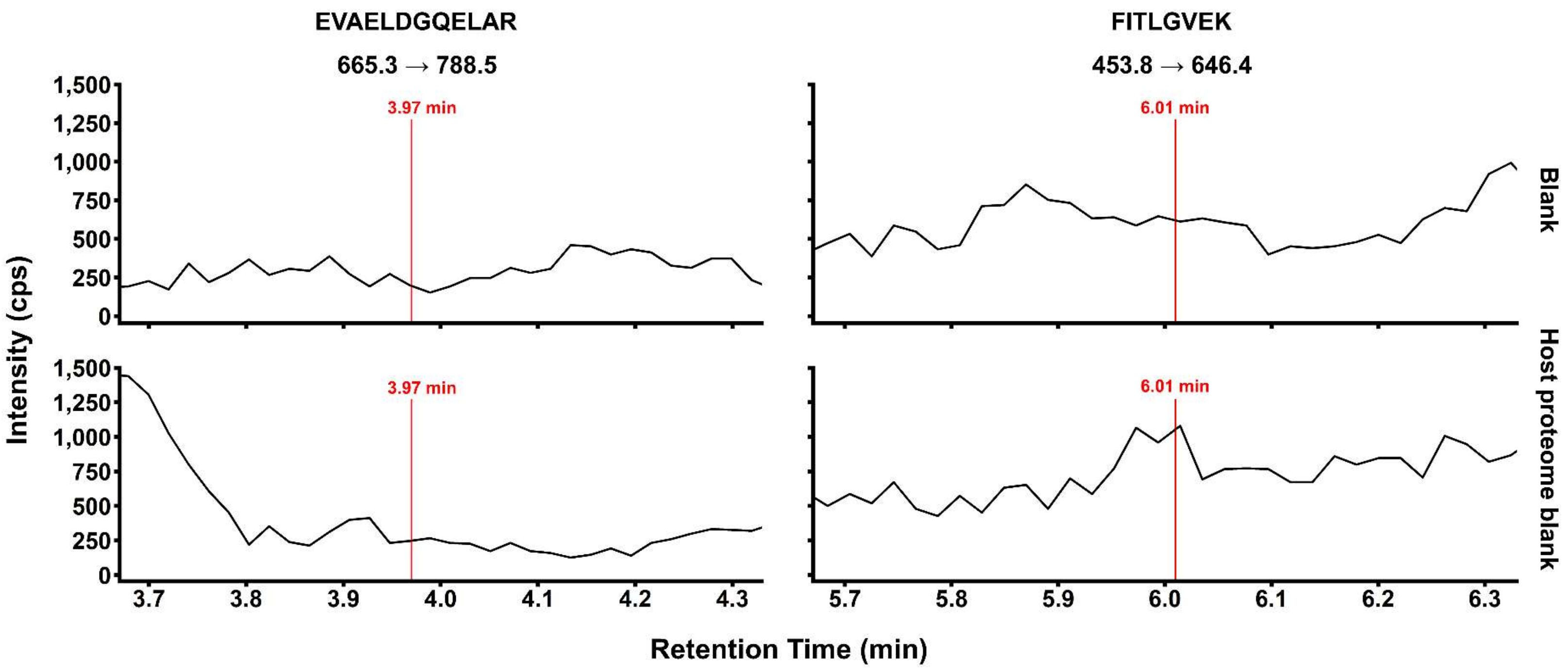

**Figure S6: MS/MS spectrum of the evolved LUZ19 peptide.** Product ion spectrum of the doubly charged precursor of EVAELDGQELAR (m/z 665.3) with annotated a, b, and y fragment ions.

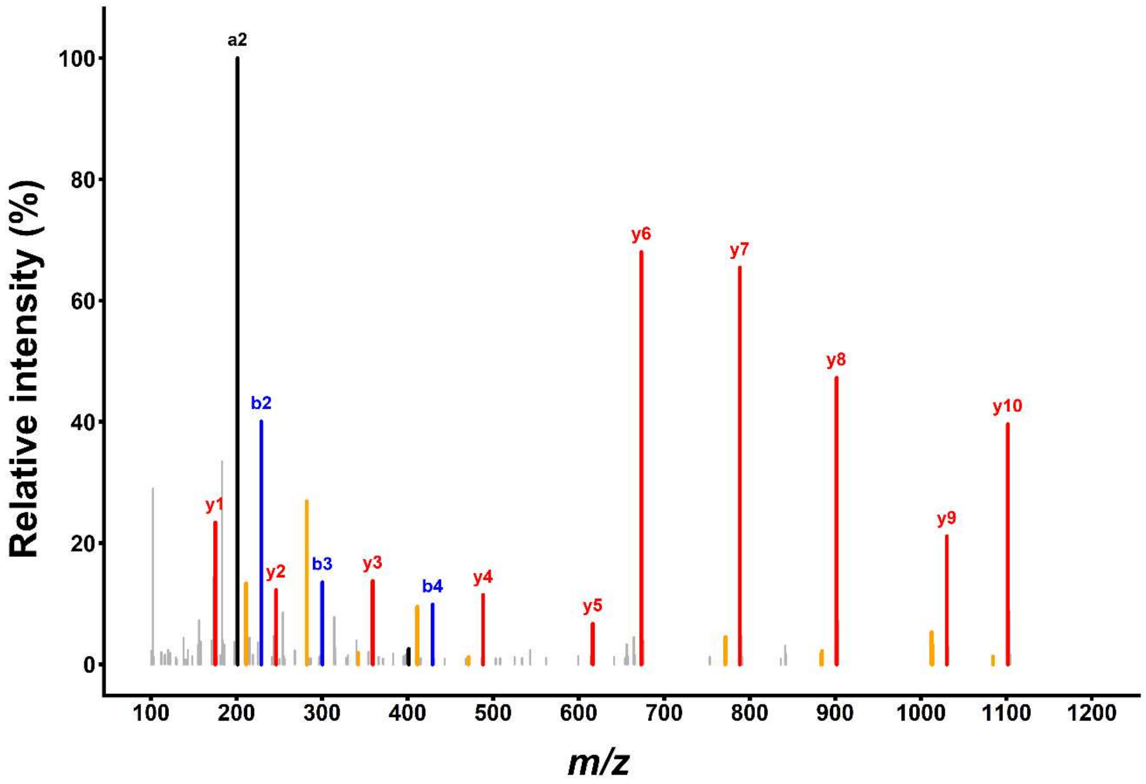

**Figure S7: Bland-Altman analysis of predicted versus observed LUZ19 phage concentrations.** Agreement between mass spectrometry-predicted and DLA-derived phage concentrations. The solid black line represents the mean difference (0.081 $\log_{10}$ units [95% CI: 0.019 to 0.144]) and the dashed red lines represent the 95% limits of agreement (−0.052 [95% CI: −0.128 to 0.024] to 0.214 [95% CI: 0.138 to 0.290] $\log_{10}$ units).

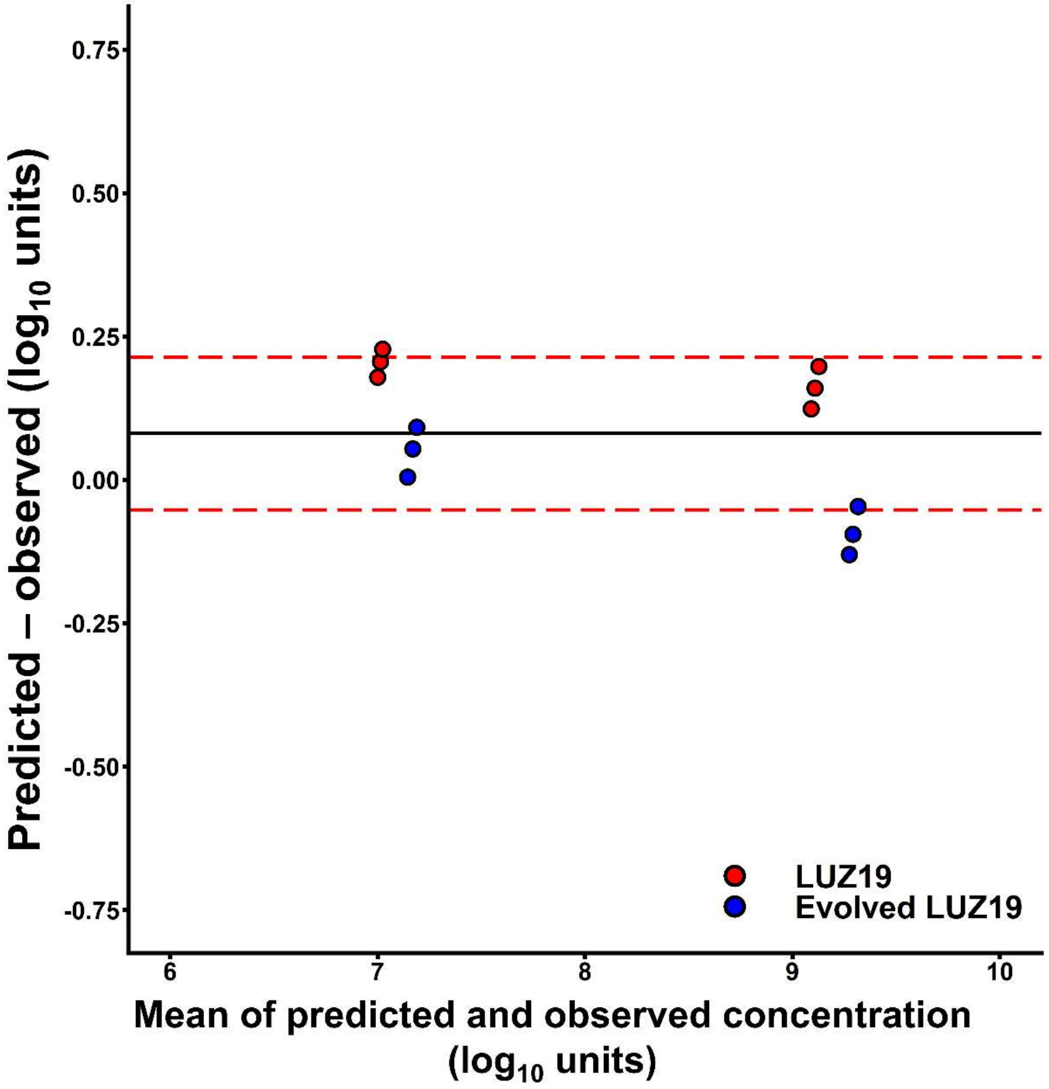

**Table S7: Efficiency of plating of LUZ19 after directed evolution.**

| Bacterial isolate | Day 0 | Day 10 | Fold-change |
|---|---|---|---|
| PAO1 | 1.00 ± 0.00 | 1.00 ± 0.00 | 1.0 |
| AR-0236 | 0.74 ± 0.10 | 1.00 ± 0.06 | 1.3 |
| AR-0240 | − | − | − |
| AR-0245 | − | − | − |
| AR-0246 | − | − | − |
| AR-0250 | − | − | − |
| AR-0254 | − | 0.00016 ± 0.00002 | − |
| AR-0257 | 0.00021 ± 0.00021 | 0.08 ± 0.02 | 390 |
| AR-0259 | − | − | − |
| AR-0270 | 1.13 ± 0.04 | 1.41 ± 0.20 | 1.3 |

*Efficiency of plating (EOP) relative to PAO1. Values are mean ± SD of three independent experiments. Host-range expansion, defined as an increase of at least one EOP class, was observed for 1 of 9 isolates (AR-0254). Fold-change is the ratio of mean EOP at Day 10 to that at Day 0.*
